# De Novo Hydration of Cryo-EM Reconstructions through Molecular Dynamics Simulations of the Excess Chemical Potential

**DOI:** 10.1101/2025.05.23.655847

**Authors:** Qinfang Sun, Sriram Aiyer, Avik Biswas, Allan Haldane, Sompriya Chatterjee, Nobuyuki Matubayasi, Dmitry Lyumkis, Ronald M Levy

## Abstract

Predicting the precise positions of water molecules at the protein interface remains a formidable challenge, fueling active research in this field. Here, we present a novel approach based on molecular dynamics simulations that utilizes statistical thermodynamic signatures of water at protein interfaces to improve the accuracy of water placement in cryo-EM maps, with apoferritin as a model benchmark system. The interaction energy of solvent with the protein is insufficient to distinguish between high- and lower-consensus water positions, consistent with reports of earlier work. Instead, we employ a detailed statistical thermodynamic analysis based on the excess chemical potential (WT) - a measure of the thermodynamic balance between the interaction energy of an interfacial water molecule with the protein and its free energy of interaction with all of the other solvent molecules. WT is proportional to the log ratio of the local density of water molecules at the protein interface to the bulk density. 85% of the top 100 water locations with the most favorable excess chemical potential values are observed in one or more high resolution Cryo-EM maps deposited in the PDB, and 70% of the top 200 water locations indexed by excess chemical potential are observed in the cryo-EM maps. This work paves the way for the development of a cryo-EM refinement tool that integrates molecular dynamics simulations with cryo-EM data for high-resolution modeling of water networks.

## Introduction

Water is critical for ligand pharmacology and can contribute significantly to ligand binding thermodynamics. For example, water displacement by a single chemical group within bound ligand can contribute substantially to gains in binding affinity, from both enthalpic and entropic considerations^1–4^. Water also plays a critical role in drug resistance. For example, displacement of waters bound within the active site of lentiviral integrases has been proposed to be the initiating step for a key mechanism of drug resistance^5^, which is conserved among lentiviral integrases^6^. Many such examples exist in literature^7–10^. Thus, the proper identification of water molecules, especially within regions bound by ligands, is crucial for pharmacological efforts to improve drug design.

The standard approach for placing waters into cryo-EM density maps includes the steps outlined below. First, a model of the macromolecular (protein or nucleic acid) assembly is built into experimentally derived cryo-EM density. This model is subsequently refined to optimize the fit into density using real space refinement tools, such as those implemented within Phenix^11,12^ or Refmac^13^. At this stage, select regions of the model will typically need to be adjusted and/or rebuilt, which can be performed within Coot^14,15^ or similar tools. The process of building/rebuilding and refinement is iterative, until the fit to density is maximized. Once a satisfactory model is derived, the next step is to place waters into residual peaks of density that are not explained by the protein or nucleic acid atoms. This can be accomplished using built-in tools within Phenix, Refmac, or Coot. There are standard considerations for assigning waters into peaks of density. The primary consideration is the shape and intensity of the density peak above a user-defined threshold for the experimental map. Water molecules generally exhibit small ovoid/spherical densities within experimental maps, which are then identified by the programs. Secondary considerations are intended to help validate whether a peak corresponds to a true water position. Since water molecules are typically found within hydrogen-bonding distance (between ∼2.5–3.5 Å) to proteins, nucleic acids, or ligand atoms, the neighboring atomic contacts, coordination geometry, and bond angles, are additional considerations to be taken into account. Waters frequently have B-factors that are similar to their surrounding atoms, and thus B-factors for assigned peaks that are unusually low may be discarded. The appearance of water peaks is also dependent on experimental map resolution. At higher resolution, there are more peaks that are observed in experimental maps, and accordingly, one can apply more stringent criteria to avoid assigning noise as a water. Below ∼3.0 Å resolution, most peaks assigned to water disappear^16,17^. (As a final step, it is essential to manually inspect each proposed water molecule to ensure it fits the density, forms plausible hydrogen bonds, does not create steric clashes, and satisfies both chemical and structural expectations. These standard criteria help distinguish real water molecules from noise, or from other small density peaks.

In addition to the considerations above, there are factors that are more specific to cryo-EM maps, which can further confound standard hydration protocols. (i) Excessive sharpening of the cryo-EM map will elevate noise, producing additional peaks that can be confused with water densities. (ii) The presence of either conformational or compositional heterogeneity within the reconstructed map (ubiquitous in cryo-EM reconstructions) will yield lower resolution densities for select regions of a map. However, standard sharpening methods based on uniformly assigned negative temperature factors^18^, which are predominantly used in the field, do not account for local resolution measures and will “over-sharpen” lower-resolution regions of a map, thus producing spurious peaks that may be misinterpreted as waters. Although numerous “local” sharpening protocols have been proposed to mitigate this problem^18–20^, they are not necessarily in standard use. (iii) Finally, there are currently no good methods to distinguish between metals or ions from waters within experimental densities in cryo-EM, although recommendations, based in part on the local chemical environment, have been discussed^16^. In X-ray crystallographic maps, methods based on anomalous scattering during X-ray data collection can yield clues and help confirm the presence and identity of specific metal ions in the density^16,21^. However, no such direct methods – based on physical principles – exist in cryo-EM, which makes distinguishing between waters and metals an acute problem. As a specific case study, in maps of lentiviral intasomes, the same peak within the active site of the ligand-bound complex was identified as a chloride ion in one structure^5^ but a water molecule in another^1^. Collectively, these issues indicate that experimental placement – and validation – of waters into cryo-EM densities remains challenging.

Several newer methods have been devised to place waters into experimental densities, to simplify the above steps, and/or to improve and validate the accuracy of placement. A Segmentation-guided Water and Ion Modeling (SWIM) protocol, which employs a semi-automated procedure for segmenting maps, establishing an appropriate map threshold, peak sorting and identification, and peak assignment based, in part, on the nearby chemical environment, was proposed to more accurately place waters^22^. Recently, the Metric Ion Classification (MIC) tool has been introduced, which passes a fingerprint representation for the neighboring chemical environment to a machine learning approach to obtain probabilities for water and ion positions within peaks of density^23^. For validation, the UnDowser functionality within Molprobity checks for all-atom contact clashes to distinguish between the diverse cases when there is something other than water assigned to a peak of density^24^. The user receives a diagnostic table of clashes, which can then be manually inspected to remove offenders. The CheckMyMetal server^25,26^, though geared toward metal ion identification, can also help to distinguish between metal ions and water molecules, based on overall structure quality and internal validation metrics. Recent advancements in the field involve the use of neural networks positioning of water molecules around proteins. HydraProt^27^ is a deep learning tool designed to predict the positions of water molecule oxygen atoms around proteins. To the best of our knowledge, a molecular dynamics-based framework has not been employed to identify and place waters into cryo-EM maps.

Solvent refinement in cryo-electron microscopy (cryo-EM) structures entails optimizing the modeling of the solvent region surrounding the macromolecule to boost the signal-to-noise ratio in density maps, both improving the precision and resolution of the final conformation^28,29^ and understanding the role of the solvent structure in the protein function. This process is important for obtaining high-resolution macromolecular models and includes several key methods: solvent masking^30^, which involves creating a mask to distinguish the solvent from the macromolecule, thus concentrating refinement efforts on areas of interest; solvent modeling^28^ which includes explicitly modeling solvents like water around the macromolecule based on the density map to enhance structural accuracy; and density modification^31^, which employs techniques such as histogram matching or solvent flattening to adjust the density in the solvent area, making the macromolecule’s features more distinct.

Accurate identification of water is still an ongoing area of research even in atomic resolution X-ray structures and is becoming an important focus in cryo-EM map analysis^32^. phenix.douse from the Phenix suite is a commonly used tool for building water molecules into cryo-EM maps^33^. Placing water molecules into high-resolution cryo-EM maps using current state-of-the-art software like Phenix can be challenging due to two main issues: distinguishing real water from noise and setting appropriate thresholds. It’s often hard to tell if the observed density in the map at a specific threshold is a water molecule or just random noise. Additionally, defining the right thresholds to differentiate the real signal from the noise in the entire cryo-EM map is complicated.

Molecular-dynamics (MD) based approaches can provide valuable information to facilitate refining waters in cryo-EM structures by offering a likelihood-based method for placing waters at key locations, validating observed densities in cryo-EM maps, and serving as an alternative when water densities are not fully distinguishable from noise, which is often the case, except at the highest resolutions. By simulating the behavior of water molecules around the macromolecule, MD can help identify probable water positions and improve the accuracy of solvent identification, and refinement in cryo-EM structures. Recently, Roh et.al integrated cryo-EM and MD simulations to cross-validate ordered water molecules along the proton path of yeast V_o_ in cryo-EM maps^34^.

In this current work, we have evaluated the excess chemical potential of water at protein interfaces, and used these signatures as a novel tool to identify the locations of high density interfacial waters for their subsequent placement into cryo-EM structures. This approach advances a method to build solvent structure in Cryo-EM maps based on a firm foundation in statistical thermodynamics. The statistical thermodynamic characterization of multiple interfacial phenomena is essential for understanding numerous biophysical and chemical events, including protein folding, ion channel gating, the aggregation of membrane proteins, and the formation of protein assemblies. It also has relevance to the energy industry for applications like electrolyte transport through pores^35–38^. According to multiple studies, analyzing the excess chemical potential (WT) is central to the problem of interpreting interfacial processes and the statistical thermodynamics of liquids^39–41^. The excess chemical potential of solvent at the protein interface is expressed as the difference between the free energy of insertion of the solvent at the interface and its insertion in the bulk far from the protein. The WT can also be determined by the ratio of the local density of water molecules around a protein to the bulk density^42^. We have recently shown that designing tighter binding ligands can be aided by knowledge of the excess chemical potential of hydrating water at protein-ligand binding sites^4^.

The target system we studied in this research is apoferritin (shown in **Figure 1A**), which plays a role in iron storage and regulation within living organisms^43^. Over the last decade, Apoferritin has evolved as one of the primary benchmark systems for developing novel methods for cryo-EM refinement. The advantages of this model system include its ease of production, biochemical stability, large size (450 kDa) that yields high scattering contrast within micrographs, structural homogeneity (few moving parts), and high octahedral symmetry that effectively multiples the number of particles by 24 symmetry-related copies. These properties mean that fewer particles are required to produce high-resolution maps, often from one or several hundred movies. The highest atomic-resolution cryo-EM structures have also been solved using apoferritin as a model specimen^44,45^.

**Figure 1.**
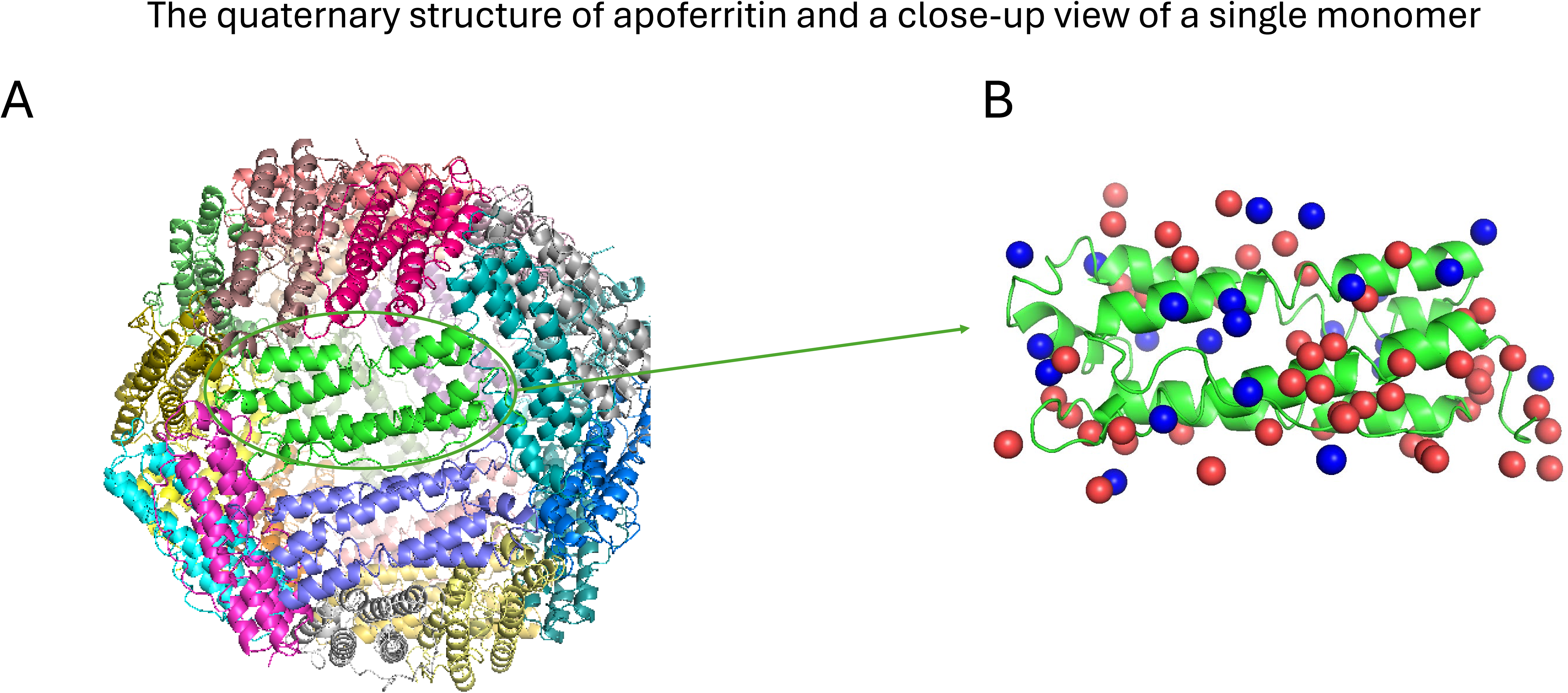
**A:** Apoferritin is a protein shell consisting of 24 subunits (7A4M). **B:** Selected high consensus(red) and lower-consensus(blue) waters are shown on a single apoferritin monomer (7A4M).

As an initial benchmarking exercise, we first compared water positions across nine high-resolution (better than 1.5 Å) experimental structures of apoferritin, identifying clusters of waters present in all nine structures, as well as clusters seen in fewer structures—ranging from one to eight. We used these water clusters to develop criteria for assigning water positions as “high consensus” (present in five or more high resolution structures) or “lower consensus” (present between one and four high resolution structures) based on the number of high-resolution cryo-EM maps that identified water at a given location. Next, we analyzed the statistical thermodynamic properties of these “high consensus” and “lower consensus” interfacial water locations by evaluating their interaction energies with apoferritin and excess chemical potentials (WT). This analysis provided insights into the correlation between the strength of the excess chemical potential and the likelihood that a water location is identified in a cryo-EM map with high or lower consensus. Finally, we investigated potential reasons for inconsistencies in the identification of water positions when comparing the experimental cryo-EM maps. We suggest how the excess chemical potential can be used to resolve these inconsistencies and speculate why some locations with highly favorable excess chemical potential are not resolved as water density in the cryo-EM maps. This study contributes to the development of a new tool for hydrating experimental reconstructions based on the principles of statistical thermodynamics that integrates MD simulations with cryo-EM data for high-resolution refinement of solvent networks. By understanding the thermodynamic properties of water, researchers can design better therapeutic molecules that exploit the unique environment at protein-water interfaces.

## Results

In this section, we first compare water positions across nine high-resolution experimental apoferritin structures. We then present the development and evaluation of a method for predicting water positions around the apoferritin structure, based on calculating the excess chemical potential (WT).

### High- and Lower-Consensus Water Positions Derived from the Comparison of Nine High Resolution Experimental Apoferritin Structures

Before analyzing the statistical thermodynamic properties of interfacial water molecules, we compared the water positions across various experimental apoferritin structures. We obtained nine high-resolution (higher than 1.5 Å ) apoferritin structures that were resolved by cryo-EM (PDB IDs: 8RQB^46^, 7A6A^44^, 8J5A^47^, 7A4M^48^, 6Z6U^44^, 7RRP^22^, 7A6B^44^, 7K3V^22^, and 7K3W^22^) (**Table S1**) from RCSB-PDB^49^. First, these nine PDB structures were aligned to identify a total of 1,626 water molecules when aggregated over all 9 PDB structures within 4 Å of a single apoferritin protomer. These positions were then grouped into “clusters” by connecting any pair of water molecules that are less than 1 Å apart from one another^22^ and identifying the disconnected sets of positions. For each cluster, the mean position of the water molecules was calculated. Clusters in which the mean position was more than 3.25 Å^50^ away from the apoferritin structure—the cutoff for the first hydration shell—were discarded, resulting in 196 unique water locations. The distribution of the number of clusters as a function of the number of water molecules per cluster is shown in **Figure 2**. Water positions identified in five or more high resolution cryo-EM structures were designated as high-consensus water positions, while those found in only one to four structures were classified as lower-consensus water positions. Selected high-consensus(red) and lower-consensus waters(blue) are shown on the surface of the apoferritin monomer (**Figure 1B**). A higher proportion of lower-consensus waters are located near the more flexible regions of apoferritin compared to the high-consensus waters.

**Figure 2:**
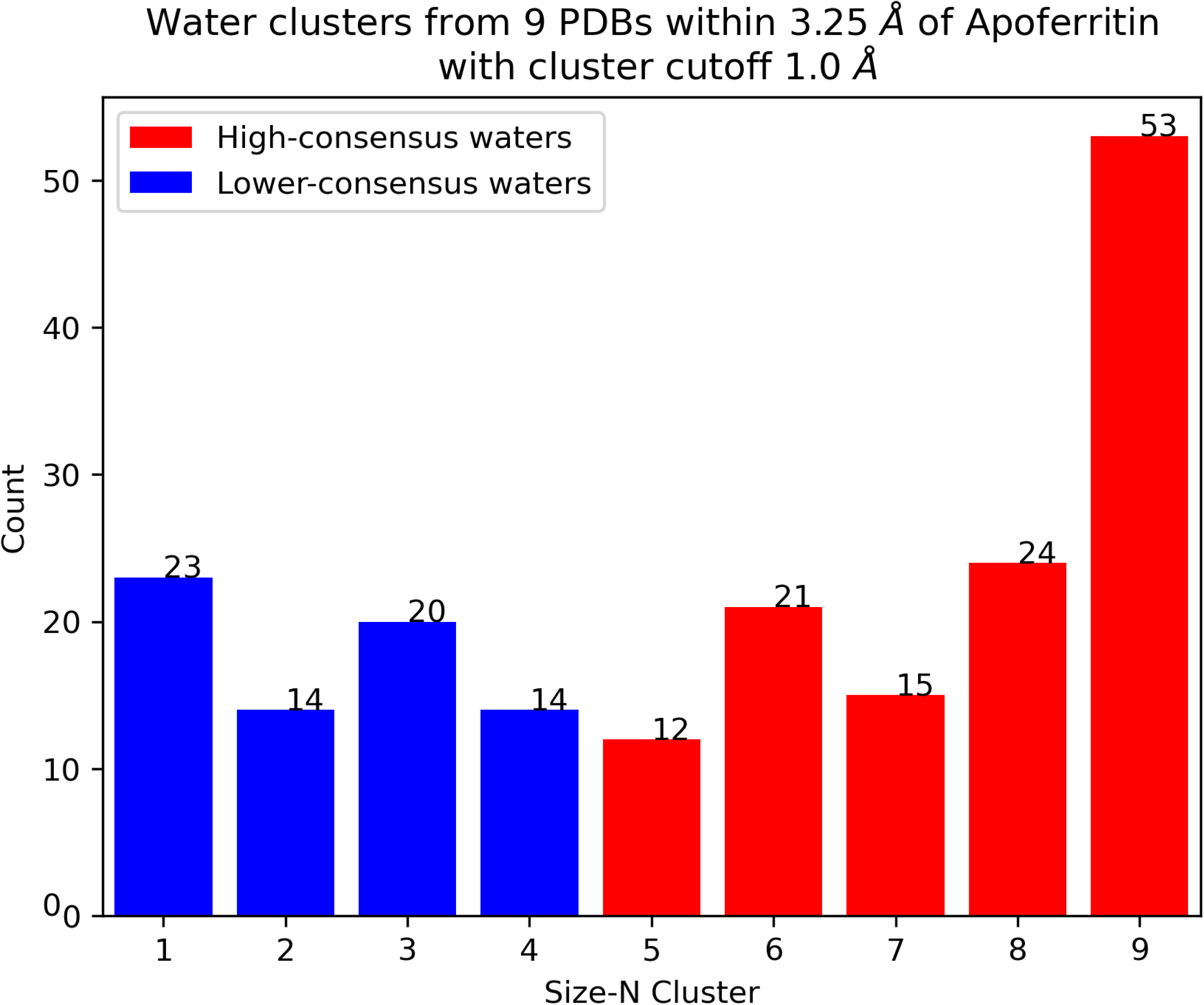
Water cluster sizes range from 1 to 9, with a size-9 cluster indicating that the water is present in all nine cryo-EM structures. The high-consensus water clusters (cluster size 5-9) are shown in red, and the lower-consensus water clusters (cluster size 1-4) are shown in blue. These water molecules are within 3.25 Å of the apoferritin structure, ensuring proximity to the protein’s surface. Additionally, they are separated by a minimum distance of 1 Å from one another to maintain spatial distinction and avoid overlapping density.

### Analysis of the Cryo-EM Apoferritin Structure using Confidence Maps at Various Resolutions

Following the identification of high-consensus and lower-consensus water positions, we conducted a comparative analysis of cryo-EM density at these water positions by reprocessing a cryo-EM density map of Apoferritin (EMDB 11638, EMPIAR-10424)^51^ and assigning a confidence score [see methods]. Most high-consensus positions that are consistently observed across all nine PDB structures exhibit a density distribution of approximately spherical shape (**Figure 3 A-D** at resolution 1.34Å). Conversely, about half of the lower consensus water positions (**Figure 3 E-H** at resolution 1.34Å) were characterized by an irregular density distribution. We also examined how the number of waters identified from the cryo-EM density changes with varying resolutions.

**Figure 3:**
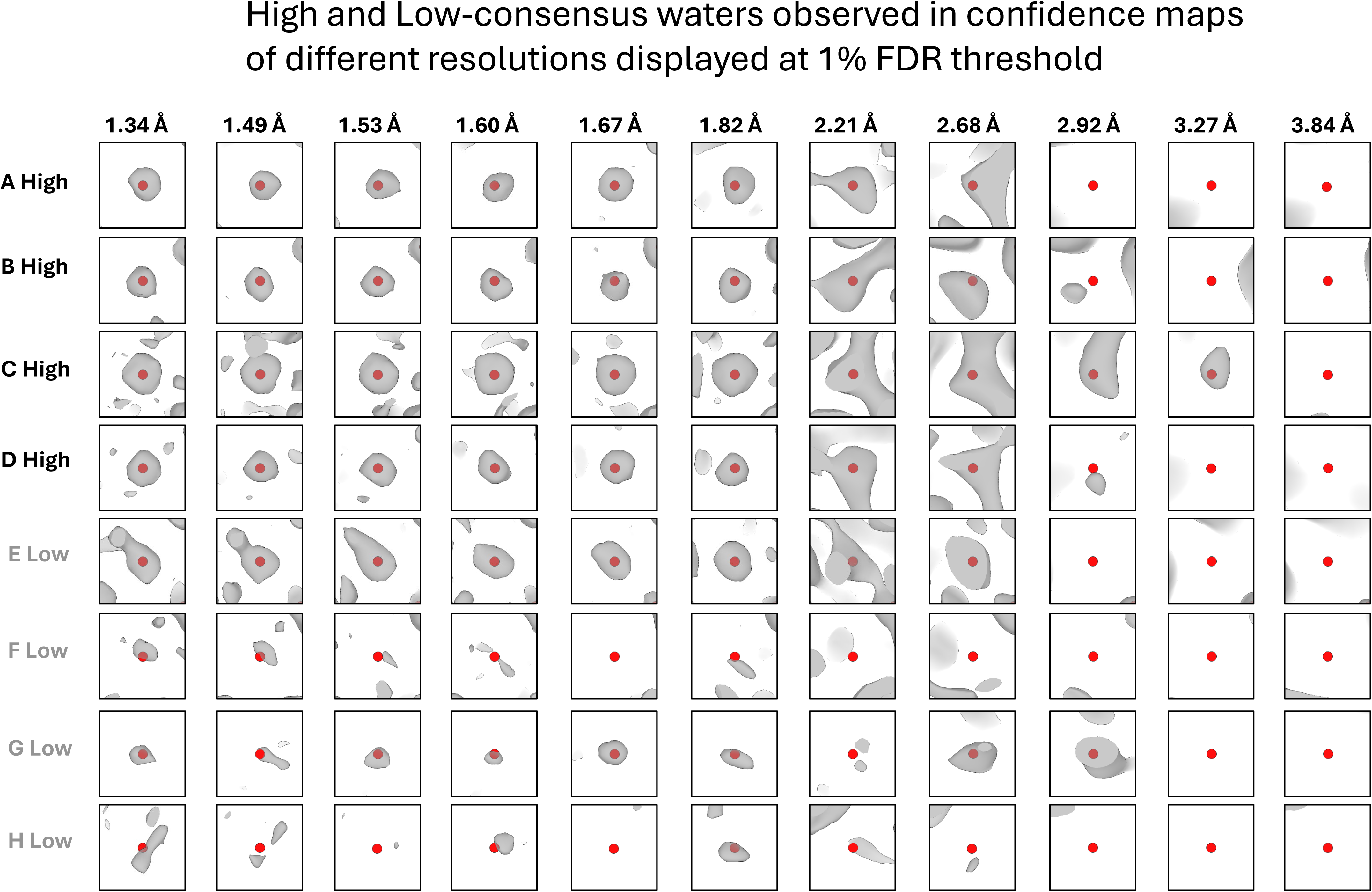
High-consensus (A–D) and low-consensus (E–H) waters are observed in confidence maps derived from cryo-EM data across a range of resolutions (1.34 Å, 1.49 Å, … 3.84 Å). These maps are displayed at a 1% false discovery rate (FDR) threshold.

To assess how the number of identified waters changes with resolution, confidence maps derived from cryo-EM density maps at various resolutions were displayed using three different false discovery rate (FDR) thresholds to determine whether water densities were unambiguously observable at the modeled coordinates. These thresholds determine the level of stringency applied when identifying water densities in the maps, with lower FDR values providing higher confidence in the observed densities. We obtained different reconstructions at progressively lower/poorer resolutions by decreasing the number of particles used in the final reconstruction of the same cryo-EM map (EMDB 11638^48^). **Figure S1** shows the cryo-EM apoferritin maps at various resolutions along with Y168 displayed at the bottom of the respective map to visualize how the map resolution varies as a function of resolution. The detailed procedure for this analysis is provided in the **Methods** section.

Unambiguous high-consensus water positions are detected even in confidence maps of lower-resolution reconstructions (**Figure 3 A-D** at resolution 2.21Å), while lower-consensus water positions exhibit greater variability across the resolution spectrum compared to high-consensus water positions (**Figure 3 E-H** at resolution 1.67Å and **Table S2**). **Figure 4** illustrates the number of observable waters per residue as a function of resolution. The inclusion of water molecules in cryo-EM models is highly dependent on the resolution of the structure. At a resolution of 2.0 Å, an average of 0.4 water molecules per residue is included, increasing to approximately 0.6 water molecules per residue at 1.5 Å resolution. Beyond a resolution of 3.27 Å, water molecules are no longer distinguishable. Notably, the number of waters in this cryo-EM apoferritin model is lower than the established rule of thumb for X-ray crystal structures, which approximates one water molecule per protein residue at 2.0 Å resolution^52^. The results using 0.1% and 5% false discovery rate (FDR) threshold are in **Figure S2**.

**Figure 4:**
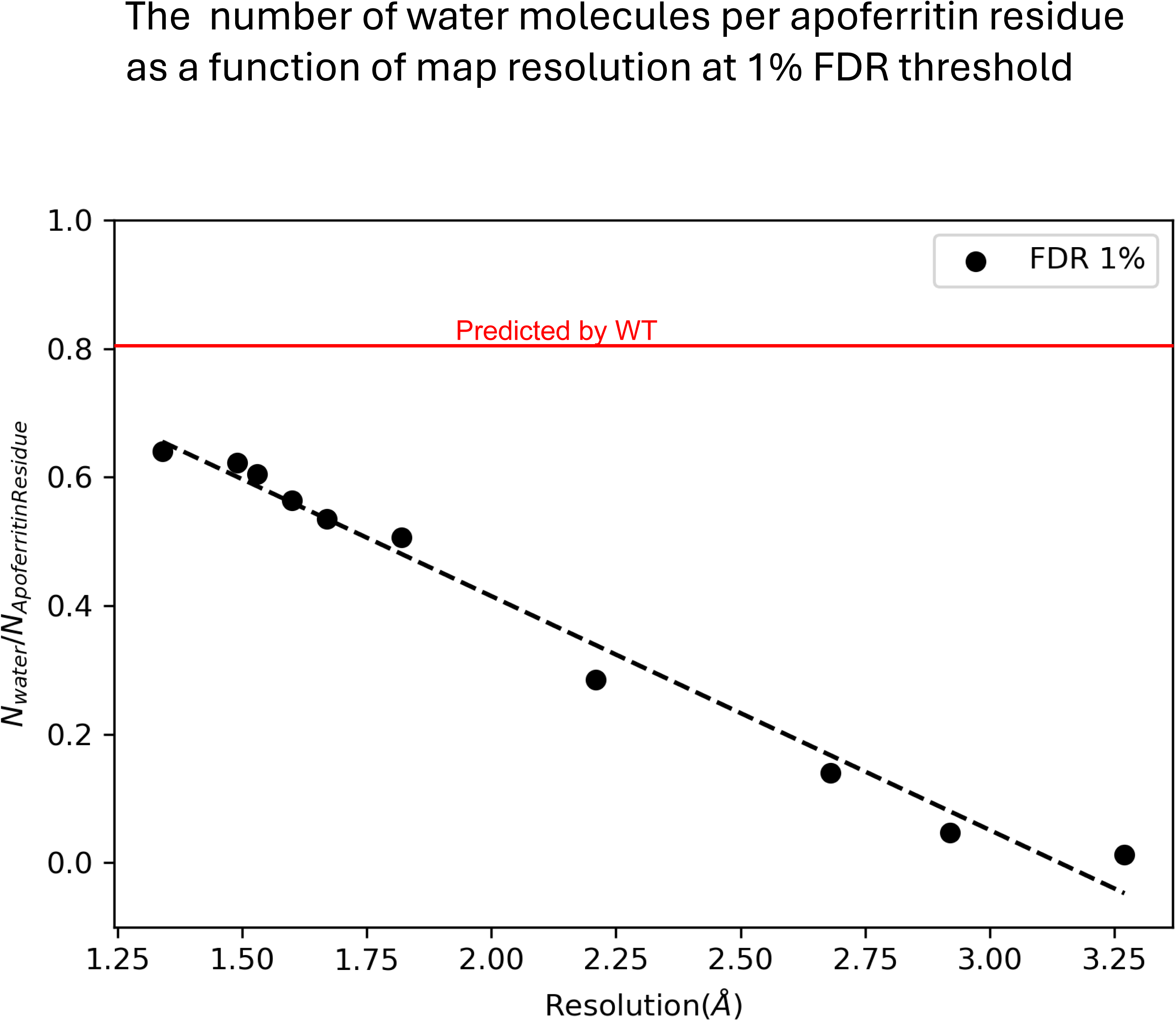
The number of water molecules per Apoferritin residue is plotted as a function of map resolution at a 1% false discovery rate (FDR) threshold.

### Statistical Thermodynamics of High- and Lower-Consensus Waters

Interfacial waters play a central role in mediating interactions between proteins and other molecules, contributing to key functional properties such as protein stability and binding. To elucidate whether the differences between high-consensus and lower-consensus water positions can be attributed to the strength of the protein-water interactions, the total interaction energy of each of these water molecules with apoferritin was calculated. As shown in **Figure 5A**, the interaction energies of the high-consensus and lower-consensus water positions overlap, i.e. they are not well separated as a function of the interaction energy with the protein, which indicates that interaction energy with the protein alone does not sufficiently distinguish between these two groups. This underscores the need for alternative approaches to more effectively differentiate between high- and lower-consensus water positions. Onuchic et al.^53^ explored the complex role of water in protein folding and function, highlighting the limitations of traditional energy calculations and emphasizing the need for more detailed thermodynamic metrics.

**Figure 5.**
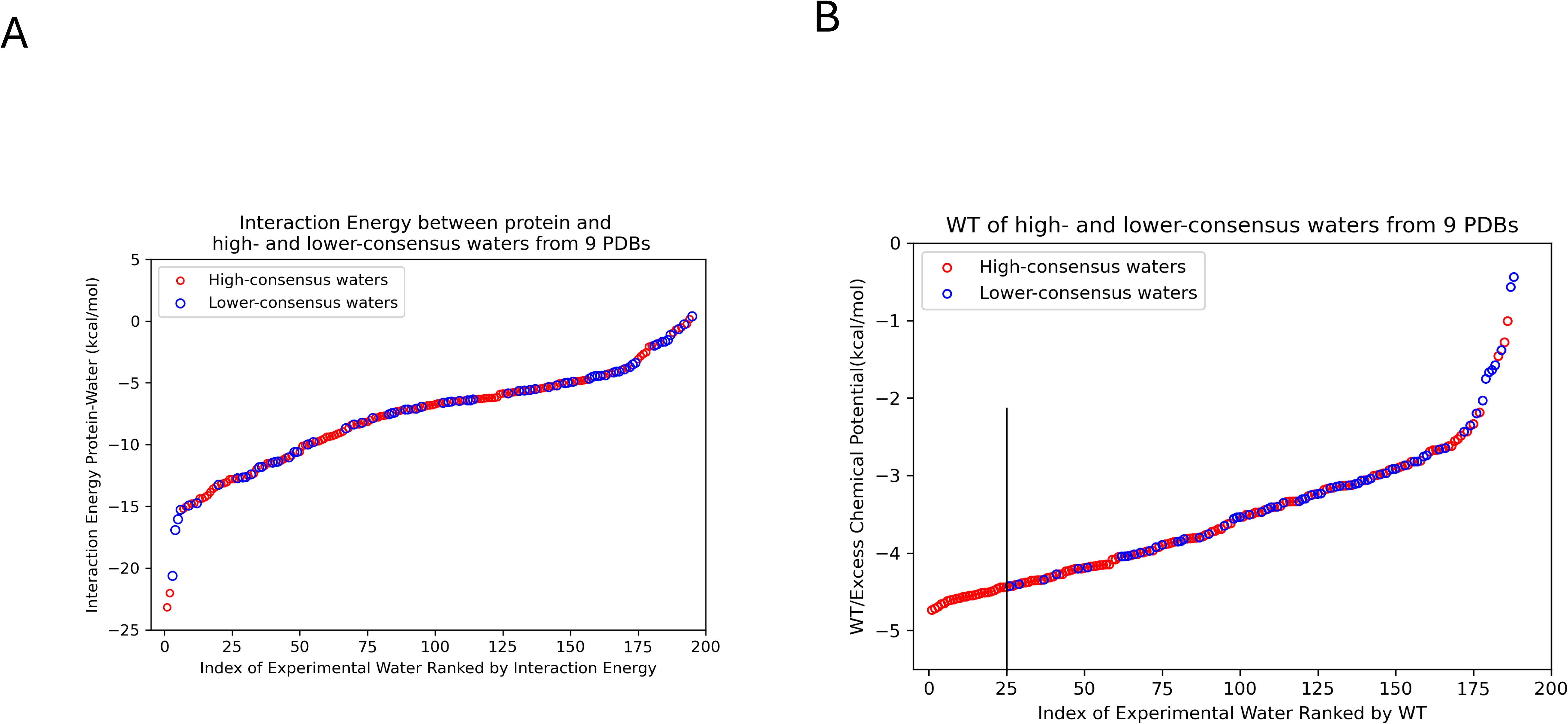
**A:** The interaction energy between the protein and selected high-consensus (red) and lower-consensus (blue) water positions derived from the 196 unique water positions is shown. **B:** It illustrates the excess chemical potential (WT) values for the same selected high-consensus (red) and lower-consensus (blue) water positions.

One such alternative method involves calculating the “excess chemical potential” (schematic diagram shown in **Figure 7**), which quantifies the extent to which the presence of the protein alters the thermodynamic properties of water molecules compared to their behavior in bulk water. The excess chemical potential for a water molecule at position (x) is a thermodynamic quantity that can be estimated by the difference between the solvation-free energy for growing a water molecule at (x), ΔF(x), and the solvation-free energy for growing a water molecule in the bulk Δ F(∞) (i.e. the pure liquid). The excess chemical potential is the “work to transfer” (WT) a water molecule from the bulk to the interfacial position at (x); it is proportional to the log ratio of the local density (ρ(x)) of water molecules at positions located at the interface with the protein to the bulk density(ρ(∞))^39^. Details of the excess chemical potential (WT) calculations by the density ratio method are provided in the methods. Our WT results, shown in **Figure 5B**, indicate that high-consensus water positions have significantly more favorable WT values compared to lower-consensus water positions. This implies that waters at high-consensus positions are more localized, and therefore the water density is higher than at lower-consensus positions, according to **Equation 1**. The WT values for high-consensus waters generally range from -3.5 to -4.8 kcal/mol, while low-census waters typically range from -1.2 to -3.5 kcal/mol. Of the 25 water locations with the most favorable WT values in **Figure 5B** (left of the black vertical line), 0 corresponds to a lower-consensus water position, while 7 out of 25 water locations with the most favorable interaction energies (left of the black vertical line in **Figure 5A**) correspond to low-consensus water positions. This distinction suggests that WT values are an effective metric for differentiating high-consensus waters from lower-consensus waters on the surface of apoferritin.

**Figure 6** illustrates the confidence scores derived from Cryo-EM confidence maps at two different resolutions: 2.7 Å and 1.3 Å. Water positions at high resolution (1.3 Å) are more likely to have a confidence score of 1, indicating that the water is clearly observed at that location in the Cryo-EM map. In contrast, a confidence score of 0 signifies the water is not observed at a specific false discovery rate (FDR = 1%; see the following section). Similarly, **Table 1** shows a strong tendency for waters to be assigned a confidence score of 1 at 1.3 Å resolution. At 2.7 Å resolution, however, waters are more frequently assigned a confidence score of 0 (63 instances) compared to a score of 1 (14 instances). These findings confirm that the number of experimentally observed water molecules strongly depends on the resolution of the Cryo-EM map.

**Figure 6:**
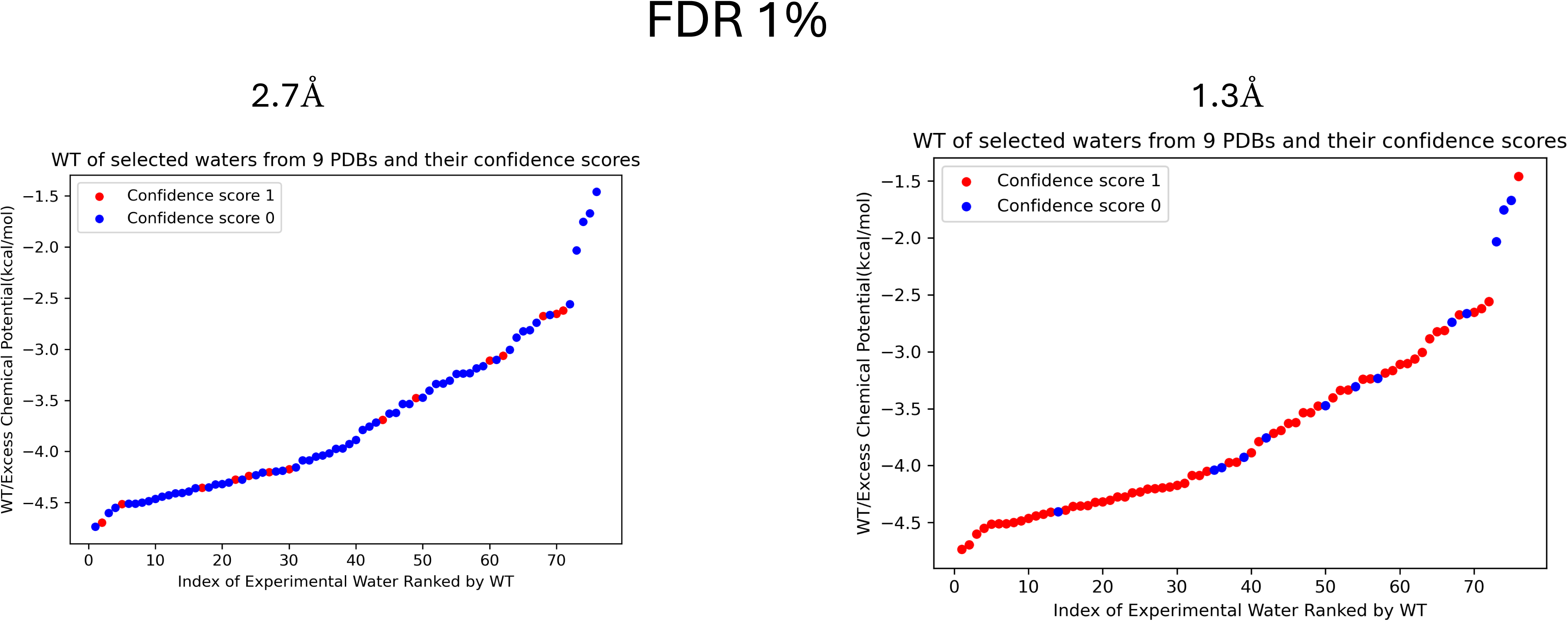
The excess chemical potential (WT) values are plotted at two different resolutions **A**:2.7 Å, **B**: 1.3 Å, with waters having a confidence score of 1 (indicating clear observability) shown in red, and those with a confidence score of 0 (indicating a lack of density) shown in blue. These results are displayed at a 1% false discovery rate (FDR) threshold.

**Figure 7:**
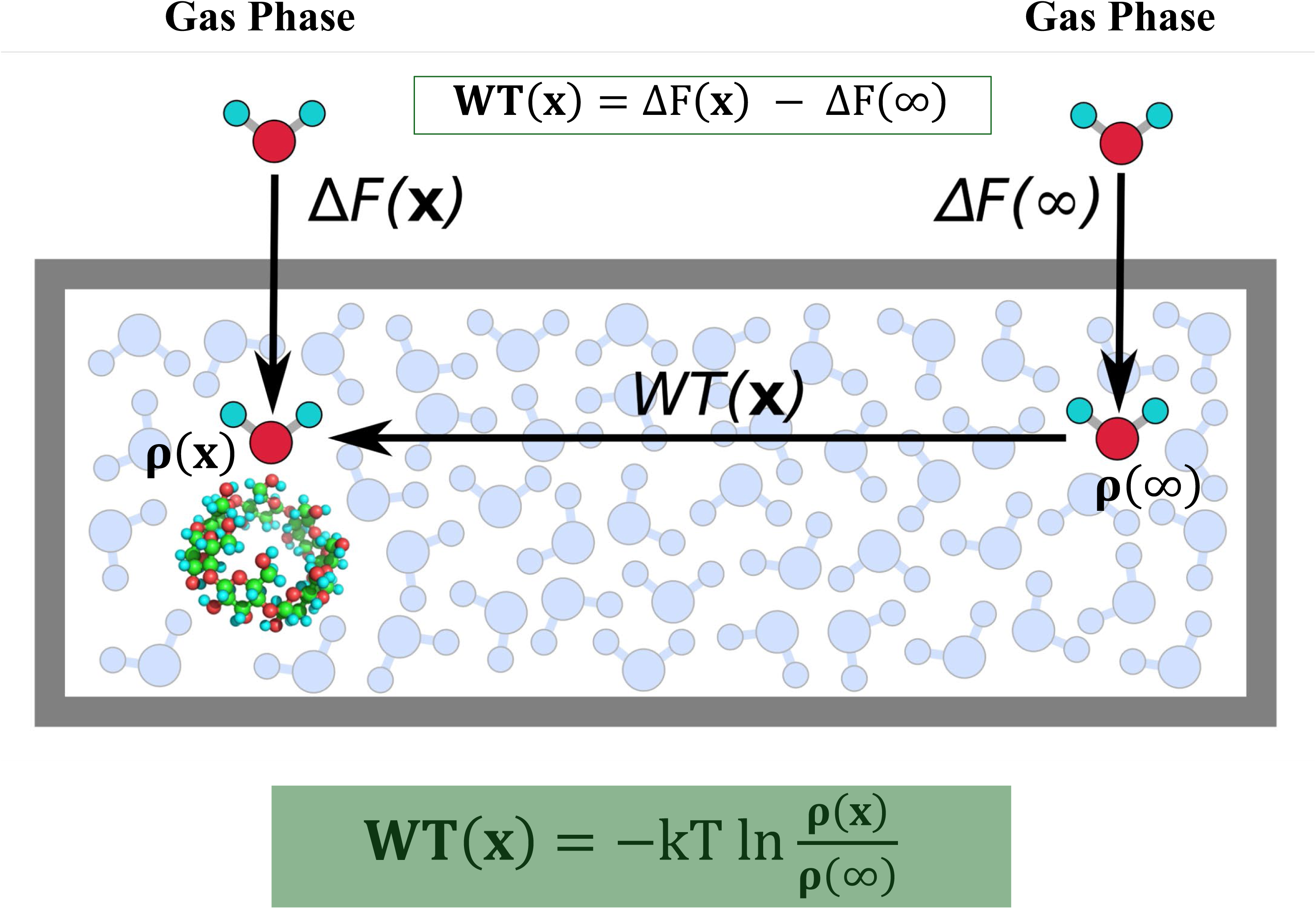
The excess chemical potential of a water molecule at x, can be estimated by the difference between the solvation-free energy for growing a water molecule at x, ΔF(x), and the solvation-free energy for growing a water molecule in the bulk ΔF(∞) (or the pure liquid ΔF(0)). The excess chemical potential can also be calculated by the log ratio of the local density (ρ(x)) of water molecules around a protein to the bulk density(ρ(∞): 0.033Å-3).

**Table 1.**
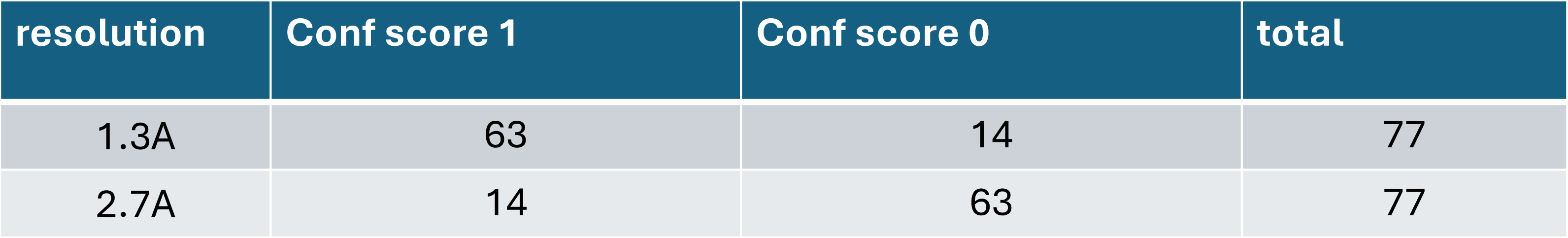

### Validation of Predicted Waters by Comparison with Experimental Waters

Despite the key role of water molecules in protein structural stability and function, accurately placing them within protein structures remains a significant challenge due to technical difficulties in resolving and modeling their precise positions^27^. To address this, we developed a novel method based on statistical mechanical principles of solvation. The excess chemical potential (WT) is a statistical thermodynamic quantity, which is proportional to the log ratio of the local density of water molecules around the protein to the bulk water density. We evaluated our method by assessing how well high- and lower-consensus water positions are reproduced by calculating the distances between the WT method-predicted water positions and those observed experimentally in apoferritin structures. These experimental structures include 9 apoferritin structures with resolutions higher than 1.5 Å (designated as group A PDBs) and 17 apoferritin structures with resolutions between 1.5 Å and 2.0 Å (designated as group B PDBs); details are provided in **Table S1**. The rationale for choosing 2.2 Å as the water match distance cutoff is twofold: first, as shown in **Figure S3**, the interaction between two hydrogen-bonded water molecules is highly repulsive at 2.2 Å, indicating that predicted and experimental waters can be considered the same when they are within this distance. Second, previous studies have demonstrated that the cutoff distance for matching experimentally determined and predicted water positions ranges from 1.4 to 2.5 Å^27,54,55^.

**Figure 8** shows that among the 200 predicted water positions based on the most favorable excess chemical potential (WT) values, 92, 29, and 21 waters are high-consensus, lower-consensus, and group B PDB water molecules, respectively (oxygen-oxygen match distance cutoff 2.2 Å). The 200 WT values range from -4.8 to -3.8 kcal/mol, corresponding to a density difference of less than 10-fold. However, the water density at these positions exceeds the bulk density by over 1000-fold. The large majority of the matched predicted water molecules are located within 1.0 Å of the experimental waters, as shown in **Figure S4**. Among the 100 water locations with the most favorable excess chemical potentials, 85% are within 2.2 A of a water molecule location identified in a Cryo-EM map of 2.0 A or higher resolution.

**Figure 8:**
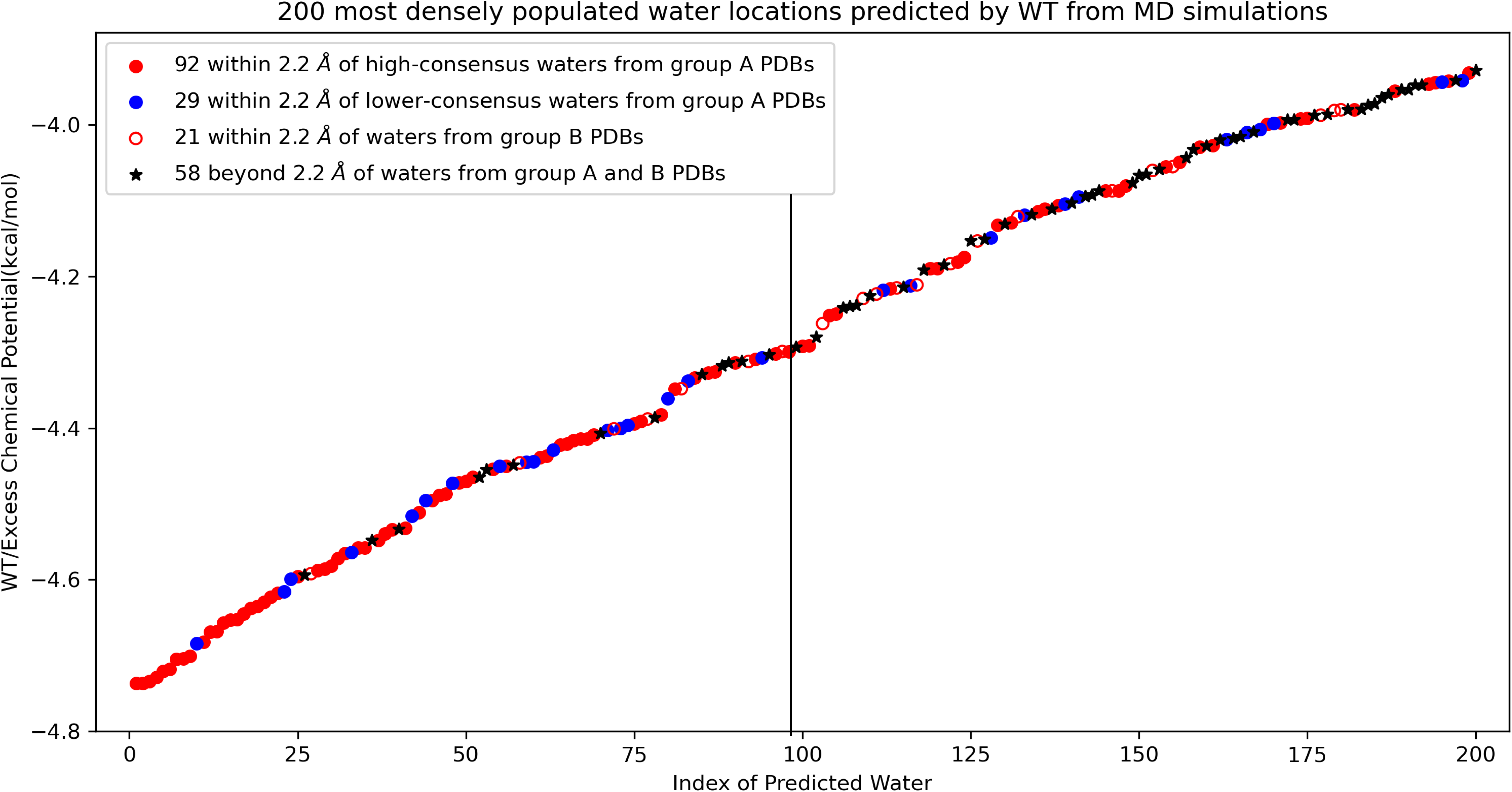
Among the 200 most densely populated water positions predicted by excess chemical potential (WT) from MD simulations, 92, 29, and 21 predicted waters are within a cutoff distance of 2.2 Å from high-consensus waters (red), lower-consensus waters (blue), and group B PDBs (red empty circle), respectively. The black star represents predicted waters that are not within 2.2 Å of any water molecules in group A or group B. Group A PDBs have resolutions higher than 1.5 Å, while group B PDBs have resolutions between 1.5 Å and 2.0 Å.

We calculated the precision and recall for predicted water positions across various excess chemical potential thresholds, as defined in the **Methods** section. **Figures 9A** and **9B** illustrate the precision and recall for three datasets: high-consensus experimental water positions from Group A PDBs (red), the sum of high-consensus and lower-consensus positions from Group A PDBs (grey), and the combined water positions from both Group A and Group B PDBs (purple).

**Figure 9:**
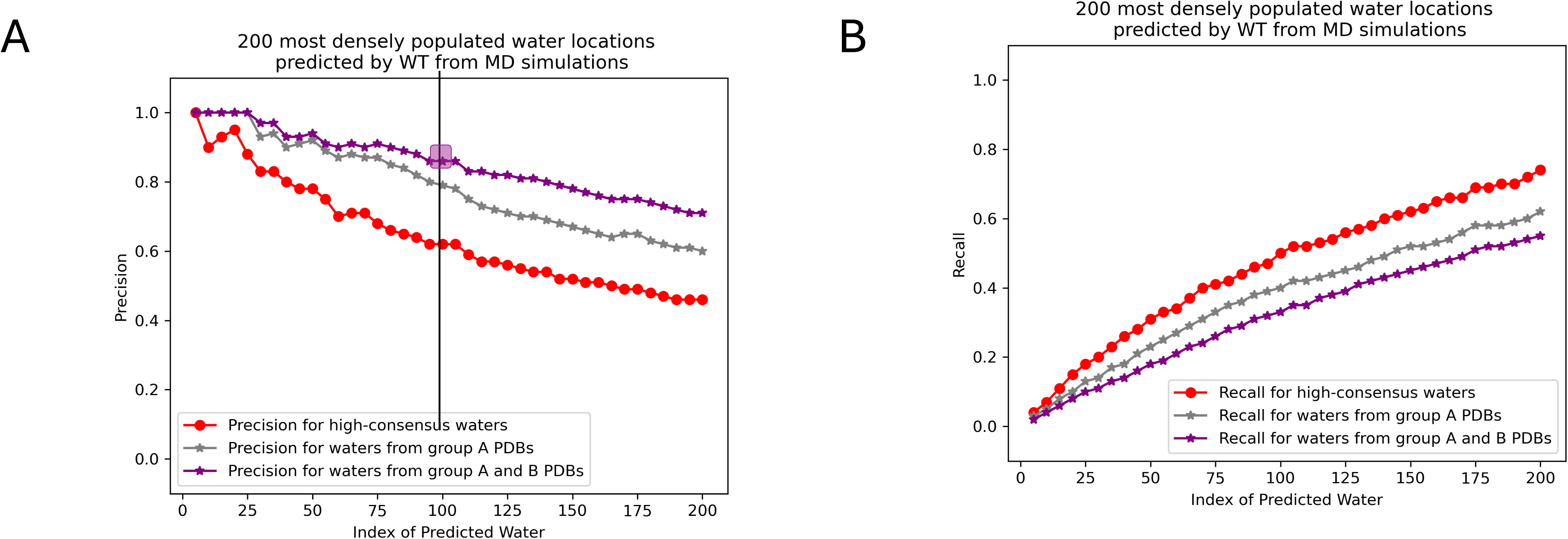
Precision (**A**) and recall (**B**) plots are shown for the predicted water positions based on WT values with a 2.2 Å distance cutoff. The curves for high-consensus waters, experimental waters from group A PDBs, and all experimental waters (group A and group B PDBs) are presented in red, grey, and purple, respectively.

In **Figure 9A**, the precision is notably higher for high-consensus waters (red curve) compared to lower-consensus waters (*difference* between the grey and red curves). The accuracy of the predicted waters is particularly striking: approximately 70% of the top 200 predicted waters, indexed by WT values, align with waters observed in experimental apoferritin structures (match distance cutoff: 2.2 Å). This precision increases to 85% when comparing the 100 most densely populated waters predicted by the excess chemical potential to experimental waters (to the left of the black vertical line in **Figure 8** and the purple square in **Figure 9A**).

Additionally, as shown in **Figure 9B**, the excess chemical potential method successfully recovers 74% of the high-consensus water positions (red curve). Using a 2.2 Å cutoff distance, 62% of the experimental waters resolved from Group A PDBs (sum of high-consensus and lower-consensus positions) are reproduced (grey curve) when a WT threshold is chosen such that the number of predicted waters closely matches the number of experimental waters (196).

Although most (>85%) of the predicted water positions with very favorable excess chemical potential values align with those observed in cryo-EM structures, a small subset of these waters does not correspond to experimentally observed water molecules. This discrepancy may be attributed to several factors. For example, some waters appear to be caged by side chains of various apoferritin residues (**Figure 10**), suggesting that the electron density associated with these waters might have been misinterpreted as part of the protein structure. Also, the pH, ion concentration, temperature, and other conditions during cryo-EM data collection differ from those used in the MD simulations. Additionally, because a single protomer is used in our MD simulations, some predicted water molecules overlap with the side chains of adjacent protomers, and waters at the interface of two or more protomers are not accurately predicted. Furthermore, in the MD simulations, the apoferritin protomer is fixed in position, which may impact the identification of waters near the flexible regions of apoferritin. The Chiu lab^34^ has also identified some reasons for the absence of MD-derived waters in experimental structures: (a) limitations of the classical force field in MD simulations, which might not fully capture polarization effects, and (b) differences in time scales between the methods—cryo-EM averages multiple chemical states over milliseconds during freezing, while MD simulations capture a single state over much shorter, nanosecond to sub-millisecond, intervals^34^. Overall, our results demonstrate that the excess chemical potential of water at the protein interface evaluated from MD simulations can be effectively used to predict water positions in cryo-EM structures.

**Figure 10:**
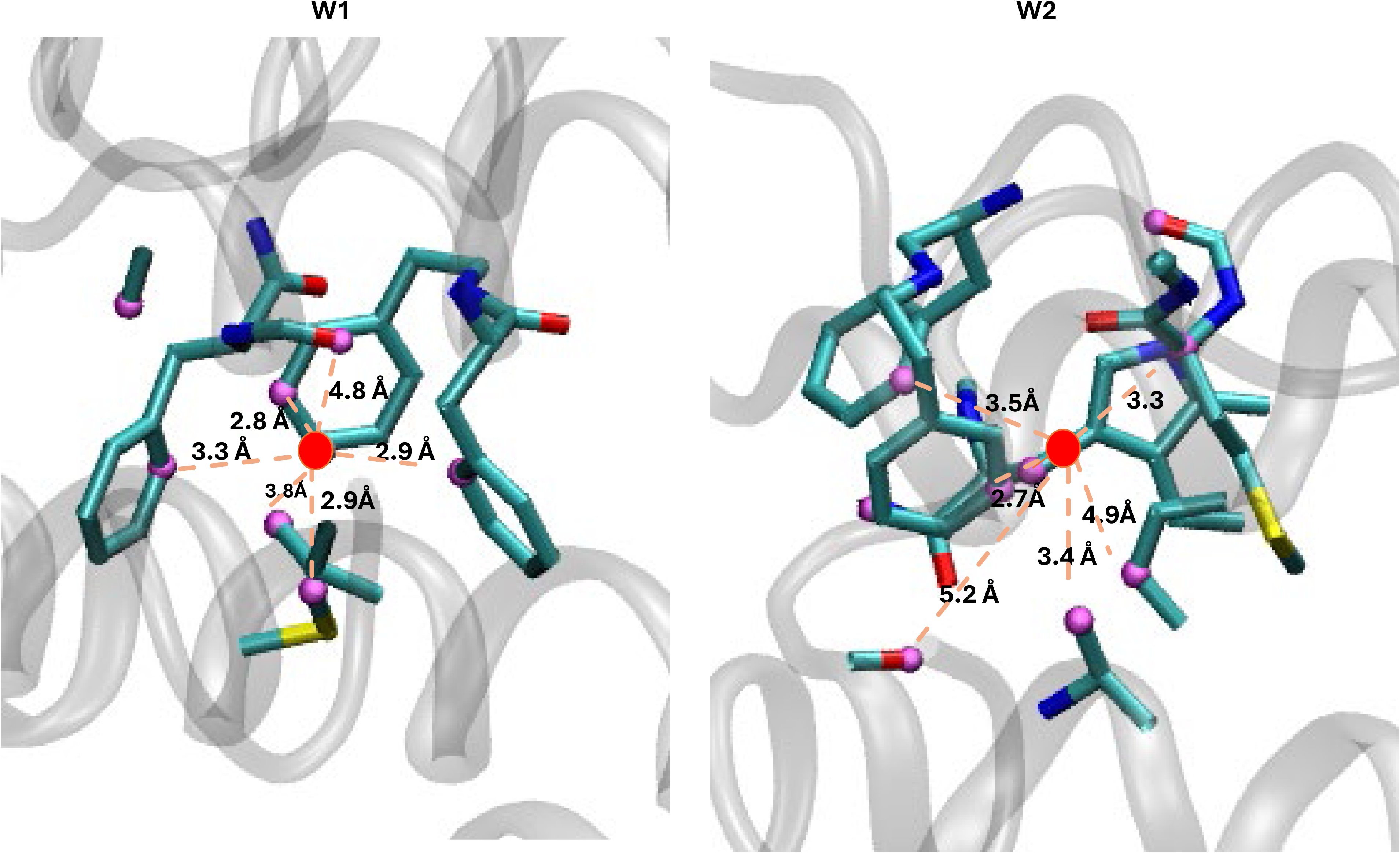
Two caged waters are shown within the apoferritin structure. Red dots represent the oxygen atoms of caged water molecules, pink indicates residue atoms within 5 Å of the caged waters, and grey cartoon represents the apoferritin structure.

## Discussion

Water assignment in cryo-EM maps remains fraught with issues. Given the challenges faced by experimentalists discussed in the introduction, it is important to develop diverse tools to assign – and validate – water positions. Although deep learning approaches can help automate water assignment, distinguish waters from metal ions or other species, and provide confidence measures based on prior data, the fundamental limitation of all current approaches is reliance on accurate training data. The often “black-box” nature of machine learning approaches, and their reliance on large, and potentially limiting training datasets, implies that physical insight into new positions may be missed. For example, a recent report highlighted how treating experimental cryo-EM maps with sharpening tools based on deep neural networks can attenuate, or completely eliminate, experimental densities corresponding to ligands and solvent^56^. In this regard, the major advantage of the current method is the ability to improve water identification within cryo-EM density maps, based on physical principles that are rooted in statistical thermodynamics.

This study demonstrates the effectiveness of using statistical thermodynamic properties of water to identify interfacial waters in cryo-EM structures of apoferritin. By comparing water positions across nine high-resolution (better than 1.5 Å) Apoferritin structures, we classified these interfacial waters into high- and lower-consensus groups depending upon the number of cryo-EM structures that included a water at that location. Using molecular dynamics simulations, we analyzed the interaction energies between high- and lower-consensus waters and the protein and examined their statistical thermodynamic properties, specifically the excess chemical potential (WT), to gain deeper insights into the structural stability of these water molecules. Our findings revealed that average interaction energy with the protein is insufficient for effectively differentiating between high- and lower-consensus water positions. However, by utilizing WT values derived from MD simulations, we are able to distinguish between high- and lower-consensus water locations. Of the 100 water positions with the most favorable excess chemical potential, ∼85% are observed in one or more apoferritin cryo-EM structures (Group A + B PDB files). Our method outperforms HydraProt^27^, which achieves a precision rate of 60%, and demonstrates significantly superior performance compared to water mesh predictions^54^, where only approximately 50% of the water molecules within the meshes were accurately identified as true positives. Also, we found a strong correlation between WT values and confidence scores derived from the confidence maps. Some waters with very favorable WT values are not observed in the experimental structures; particularly waters that appear to be “caged” by surrounding aromatic side chains. Further analysis and experimentation with variation of density thresholding may be helpful in resolving these discrepancies.

**Figure 4** illustrates that the number of observable waters in cryo-EM models strongly depends on the structure’s resolution. At resolutions approaching 3 Å, and particularly beyond ∼3.2 Å (X-ray) or ∼3.5 Å (cryo-EM), it becomes increasingly challenging to identify water densities de novo. At these lower resolutions, water features may either blend with noise or become entirely indistinguishable. For resolutions typical of most cryo-EM structures (∼2.5–3.5 Å), water densities are often difficult to resolve. As shown in **Figure 6**, waters that are clearly visible at 1.3 Å largely disappear or become too noisy to distinguish at 2.7 Å. In such scenarios, our WT-method provides a significant advantage by accurately predicting water positions and distinguishing water densities from noise. This approach not only aids in modeling but also accelerates the placement of waters in low-resolution cryo-EM structures, offering valuable insights into structural interpretation. Furthermore, we note that our MD simulations are performed without experimental density restraints, and yet the water positions with favorable WT values correlate strongly to their corresponding position within experimentally defined maps. This indicates that such tools would accordingly be useful to boost confidence in annotated water positions, and to identify *new* water positions, based on physical principles and thermodynamic signatures. The ideas are also extendable to identifying and characterizing metal ions within experimental maps and correlated networks of waters that are associated with allostery. Future approaches will build upon the current framework but will also include considerations of local resolution, density restraints to guide the MD simulations, and iterative model building and density-guided refinement approaches, for optimally hydrating experimental cryo-EM maps.

## Methods

### System Preparation and Molecular Dynamics Simulations

High-resolution cryo-EM structures serve as a foundation for identifying water molecules at protein interfaces. The starting structure of apoferritin for the MD simulation is derived from the cryo-EM structure (PDB 7A4M^48^ (chain A) shown in **Figure 1B**). The MD simulations were conducted using the GROMACS 2020.3^57^ suite. The ff99SB force field^58^ together with the explicit TIP3P solvation model^59^ was adopted to describe the protein and the solvent effects. Apoferritin was placed at the center of a cubic box and the distance between apoferritin and the edge of the box was set to be ≥ 1.0 nm. The final simulation box contains apoferritin, 23,038 water molecules and 5 Na^+^ ions, with a total of 71,892 atoms in the system. The procedure began with energy minimization to relax the initial configuration, followed by a 100 ps equilibration in the NVT ensemble and a 5 ns equilibration in the NPT ensemble, both with restraints on the protein heavy atoms. The production run was carried out in the NPT ensemble with all protein atoms fixed for 1 µs at a temperature of 300 K and a pressure of 1 atm. Temperature control was achieved using a modified Berendsen thermostat^60^, while pressure was regulated to 1 bar with a compressibility of 4.5 × 10⁻⁵ bar using the Parrinello-Rahman barostat^61^. Electrostatic interactions were treated using the particle mesh Ewald (PME)^62^ method with a real-space cutoff of 1.0 nm. The MD simulations were executed with a time step of 1 fs, and trajectory files were saved every 0.2 ps. For the WT analysis, 100,000 frames were extracted from the 1 µs trajectory by selecting every 5^th^ frame within the 300 ns to 400 ns time window.

### Interaction Energy Calculations for High- and Lower-Consensus Waters

The methodology for categorizing high- and low-consensus water positions (196 water clusters) is detailed in the **Results** section. We firstly incorporated 196 water molecules into a single apoferritin monomer (7A4M^48^). The process began with energy minimization to relax the initial configuration, followed by a 100 ps equilibration in the NVT ensemble and a subsequent 100 ps equilibration in the NPT ensemble. Before energy minimization, the system was neutralized by adding Na+ ions. During the minimization step, the 196 water oxygens were harmonically restrained, while both the 196 water oxygens and protein heavy atoms were restrained during the NVT and NPT equilibration simulations. The rest of the parameters are the same as the **System Preparation** section. Lastly, the total interaction energy between the protein and each water was calculated using the final frame of the NPT equilibration and extracted by the gmx_energy module. Nonbonded interaction energies (Coulombic and Lennard-Jones) were computed to quantify the strength of interactions between the protein and each water.

### WT calculations for High- and Low-Consensus Waters

*Note: This method cannot distinguish buried water from water on the surface of the protein. The interaction energy between water and the protein alone cannot determine the water density around the protein, which is why the WT method was developed. Also we represent the water molecules solely by the oxygen atom*.

The WT for the 196 unique water positions in **Figure 5** is calculated as described here. In **Figure 7** and **Equation 1**, the excess chemical potential of a water molecule can be estimated as the free energy difference ΔF(**x**) − ΔF(∞) or the density ratio 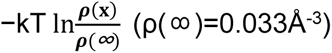

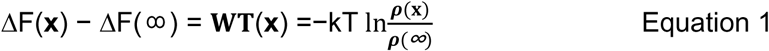

Numerically, we use 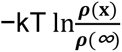 to determine WT(dV) by gridding space into small cubic voxels of volume for which we compute WT(V). When the volume approaches infinitesimally small dimensions, it can be considered as a point. In this case, it is evident that the two sides of Equation 1 are equal.

For calculating WT for the 196 water positions, we create a grid around each reference oxygen position, extending 0.5 Å in the x, y, and z directions. The grid consists of fine cubic voxels with an edge length of a=0.005 Å and volume *dV*=a^3^. Subsequently, we selectively compute WT using the densest fine voxels, that is the top 1% of the 8 million cubic voxels for each reference position, or n=80,000 fine voxels. The Boltzmann average WT is calculated as the sum of the exponential of all WT(dV) values (**Equation 1**) divided by the sum of volume of the n fine voxels (**Equation 2**).

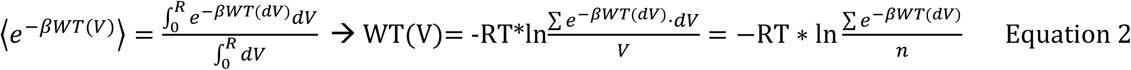

### Waters Predicted by the Excess Chemical Potential

We first search for locations of high-water density by counting all water molecules within 3.25 Å of apoferritin^50^, covering the first hydration shell, from 100,000 frames of MD simulations (see **methods section 1)** using a coarser grid with voxel edge length of 0.1 Å, resulting in a total of approximately 240 million coarse voxels. This grid is also used to visualize the water distribution around apoferritin. A sliding cube of 10 × 10 × 10 such voxels, with a total edge length of 1 Å, is used as a convolution window to identify the 1 Å sized regions with the most water molecules. We then filter these so that each 1 Å cube must be at least 2.25 Å apart from each other and at least 2.25 Å away from apoferritin. From 100,000 frames of MD trajectory, the average position of the waters within each qualifying 1 Å cube is calculated and set as the predicted water position. The center of each qualifying 1 Å cube is used as the reference position for the subsequent WT calculations.

### Evaluation Metrics: Precision and Recall

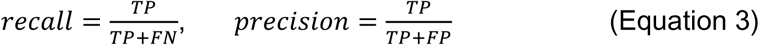

True positive (TP) predictions refer to the number of predicted water molecules that match experimental water positions in apoferritin within 2.2 Å. False positive (FP) predictions represent the number of the predicted water molecules that do not match any experimental water, while false negative (FN) predictions indicate the number of experimental water molecules that are not matched to any predicted water. In this evaluation, each predicted water is assigned to its closest experimental water, with only one prediction counted per experimental water.

### Cryo-EM Image Processing by Confidence Maps

Obtained raw movie frames from EMPIAR deposition 10424. Used 1689 movies for the entire analysis and all the data processing steps were carried out in CryoSparc v4.6^63,64^. Movies were imported using an up-sampling factor of 2 and frames were grouped into 31 fractions. Movie frames were aligned using Patch Motion Correction with an output crop factor of 1/2 and B-factor of 500 for global alignment. Micrograph CTF estimation was performed using Patch CTF estimation. Particle picking was first performed using a combination Blob Picker followed by Template Picker with suitable classes selected after 2-D classification extracted particles. All particles were then extracted using a box size of 512 pixels. All 2-D classifications have an inner and outer diameter of 125 and 140 Å. A total of 186,280 particles were obtained after iterative 2-D classification steps. An initial volume was generated through Ab Initio reconstruction of 158,100 particles with window inner and outer radius of 0.5 and 0.9 respectively and octahedral symmetry imposed. Using this initial volume, Euler angles were refined for the 186,280 using Homogeneous Refinement with iterative refinements accounting for per-particle defocus optimization, per-group CTF parameters accounting for tilt, trefoil, spherical aberration, tetrafoil and magnification anisotropy. A final refinement was performed accounting for Ewald Sphere curvature correction using positive curvature. Refined particles, volume and mask were then used for Reference-Based Motion Correction (RBMC) to optimize motion-correction of particle images and for improved dose-weighting^65,66^ . Finally, unbinned particles obtained from RBMC were re-refined to obtain a map at 1.33 Å global resolution. Random subsets of 180,000, 90,000, 45,000, 22,500, 11,250, 6,000, 3,000, 1,500, 750 and 200 particles were selected using the Particle Subsets tool and subjected to Homogeneous reconstructions to obtain maps of progressively poorer global resolutions as a function of particle number. Maps obtained from these reconstructions were used as inputs to derive Confidence Maps within the CCPEM software suite^67,68^ . A noise box of 192 pixels was used with an input pixel size of 0.228 and a box size of 1024. Correction method of FDR-BY was employed using a two-sided test procedure. Rest of the parameters were default. Positions of waters were assessed at 0.1, 1 and 5% FDR. All visualizations were performed in Chimera or ChimeraX (UCSF)^69,70^ .

## Acknowledgements

This study is supported by the National Institutes of Health R01 AI178849 (to R.M.L. and D.L.). A.B. was supported by the Eric & Wendy Schmidt AI-in-Science Postdoctoral Fellowship, a program of Schmidt Sciences. Explicit solvent MD simulations were run on the local computing resource, the CB2RR HPC cluster at Temple University, and the Expanse clusters of ACCESS (MCB100145). The WT calculations were run on the CB2RR HPC cluster using custom Python scripts.

## References

1 Passos, D. O. et al. Structural basis for strand-transfer inhibitor binding to HIV intasomes. Science 367, 810–814, doi:10.1126/science.aay8015 (2020).

2 Schiebel, J. et al. Intriguing role of water in protein-ligand binding studied by neutron crystallography on trypsin complexes. Nat Commun 9, 3559, doi:10.1038/s41467-018-05769-2 (2018).

3 Biela, A. et al. Ligand binding stepwise disrupts water network in thrombin: enthalpic and entropic changes reveal classical hydrophobic effect. J Med Chem 55, 6094–6110, doi:10.1021/jm300337q (2012).

4 Cui, D., Zhang, B. W., Matubayasi, N. & Levy, R. M. The Role of Interfacial Water in Protein-Ligand Binding: Insights from the Indirect Solvent Mediated Potential of Mean Force. Journal of chemical theory and computation 14, 512–526, doi:10.1021/acs.jctc.7b01076 (2018).

5 Cook, N. J. et al. Structural basis of second-generation HIV integrase inhibitor action and viral resistance. Science 367, 806–810, doi:10.1126/science.aay4919 (2020).

6 Li, M. et al. Mechanisms of HIV-1 integrase resistance to dolutegravir and potent inhibition of drug-resistant variants. Sci Adv 9, eadg5953, doi:10.1126/sciadv.adg5953 (2023).

7 Poornima, C. S. & Dean, P. M. Hydration in drug design. 1. Multiple hydrogen-bonding features of water molecules in mediating protein-ligand interactions. Journal of Computer-Aided Molecular Design 9, 500–512, doi:10.1007/bf00124321 (1995).

8 Wong, S. E. & Lightstone, F. C. Accounting for water molecules in drug design. Expert Opinion on Drug Discovery 6, 65–74, doi:10.1517/17460441.2011.534452 (2010).

9 Laage, D., Elsaesser, T. & Hynes, J. T. Water Dynamics in the Hydration Shells of Biomolecules. Chemical reviews 117, 10694–10725, doi:10.1021/acs.chemrev.6b00765 (2017).

10 Levy, Y. & Onuchic, J. N. WATER MEDIATION IN PROTEIN FOLDING AND MOLECULAR RECOGNITION. Annual Review of Biophysics and Biomolecular Structure 35, 389–415, doi:10.1146/annurev.biophys.35.040405.102134 (2006).

11 Adams, P. D. et al. PHENIX: a comprehensive Python-based system for macromolecular structure solution. Acta Crystallogr D Biol Crystallogr 66, 213–221, doi:10.1107/S0907444909052925 (2010).

12 Afonine, P. V. et al. Real-space refinement in PHENIX for cryo-EM and crystallography. Acta Crystallogr D Struct Biol 74, 531–544, doi:10.1107/S2059798318006551 (2018).

13 Brown, A. et al. Tools for macromolecular model building and refinement into electron cryo-microscopy reconstructions. Acta Crystallogr D Biol Crystallogr 71, 136–153, doi:10.1107/S1399004714021683 (2015).

14 Casanal, A., Lohkamp, B. & Emsley, P. Current developments in Coot for macromolecular model building of Electron Cryo-microscopy and Crystallographic Data. Protein Sci 29, 1069–1078, doi:10.1002/pro.3791 (2020).

15 Emsley, P., Lohkamp, B., Scott, W. G. & Cowtan, K. Features and development of Coot. Acta Crystallogr D Biol Crystallogr 66, 486–501, doi:10.1107/S0907444910007493 (2010).

16 Richardson, J. S. et al. Model validation: local diagnosis, correction and when to quit. Acta Crystallogr D Struct Biol 74, 132–142, doi:10.1107/S2059798317009834 (2018).

17 Pintilie, G. et al. Measurement of atom resolvability in cryo-EM maps with Q-scores. Nat Methods 17, 328–334, doi:10.1038/s41592-020-0731-1 (2020).

18 Rosenthal, P. B. & Henderson, R. Optimal determination of particle orientation, absolute hand, and contrast loss in single-particle electron cryomicroscopy. J Mol Biol 333, 721–745, doi:10.1016/j.jmb.2003.07.013 (2003).

19 Terwilliger, T. C., Sobolev, O. V., Afonine, P. V. & Adams, P. D. Automated map sharpening by maximization of detail and connectivity. Acta Crystallogr D Struct Biol 74, 545–559, doi:10.1107/S2059798318004655 (2018).

20 Ramirez-Aportela, E. et al. Automatic local resolution-based sharpening of cryo-EM maps. Bioinformatics 36, 765–772, doi:10.1093/bioinformatics/btz671 (2020).

21 El Omari, K., et al. Utilizing anomalous signals for element identification in macromolecular crystallography. Acta Crystallogr D Struct Biol 80, 713–721, doi:10.1107/S2059798324008659 (2024).

22 Zhang, K., Pintilie, G. D., Li, S., Schmid, M. F. & Chiu, W. Resolving individual atoms of protein complex by cryo-electron microscopy. Cell Res 30, 1136–1139, doi:10.1038/s41422-020-00432-2 (2020).

23 Shub, L., Liu, W., Skiniotis, G., Keiser, M. J. & Robertson, M. J. Metric Ion Classification (MIC): A deep learning tool for assigning ions and waters in cryo-EM and x-ray crystallography structures. bioRxiv, 2024.2003.2018.585639, doi:10.1101/2024.03.18.585639 (2024).

24 Prisant, M. G., Williams, C. J., Chen, V. B., Richardson, J. S. & Richardson, D. C. New tools in MolProbity validation: CaBLAM for CryoEM backbone, UnDowser to rethink “waters,” and NGL Viewer to recapture online 3D graphics. Protein Sci 29, 315–329, doi:10.1002/pro.3786 (2020).

25 Gucwa, M. et al. CMM-An enhanced platform for interactive validation of metal binding sites. Protein Sci 32, e4525, doi:10.1002/pro.4525 (2023).

26 Zheng, H. et al. Validation of metal-binding sites in macromolecular structures with the CheckMyMetal web server. Nat Protoc 9, 156–170, doi:10.1038/nprot.2013.172 (2014).

27 Zamanos, A., Ioannakis, G. & Emiris, I. Z. HydraProt: A New Deep Learning Tool for Fast and Accurate Prediction of Water Molecule Positions for Protein Structures. Journal of Chemical Information and Modeling 64, 2594–2611, doi:10.1021/acs.jcim.3c01559 (2024).

28 Shang, Z. & Sigworth, F. J. Hydration-layer models for cryo-EM image simulation. Journal of structural biology 180, 10–16, doi:10.1016/j.jsb.2012.04.021 (2012).

29 Moulinier, L., Case, D. A. & Simonson, T. Reintroducing electrostatics into protein X-ray structure refinement: bulk solvent treated as a dielectric continuum. Acta Crystallographica Section D Biological Crystallography 59, 2094–2103, doi:10.1107/s090744490301833x (2003).

30 Serna, M. Hands on Methods for High Resolution Cryo-Electron Microscopy Structures of Heterogeneous Macromolecular Complexes. Frontiers in molecular biosciences 6, 33–33, doi:10.3389/fmolb.2019.00033 (2019).

31 Cowtan, K. D. & Zhang, K. Y. J. Density modification for macromolecular phase improvement. Progress in Biophysics and Molecular Biology 72, 245–270, doi:10.1016/s0079-6107(99)00008-5 (1999).

32 Zhang, K., Pintilie, G. D., Li, S., Schmid, M. F. & Chiu, W. Resolving individual atoms of protein complex by cryo-electron microscopy. Cell research 30, 1136–1139, doi:10.1038/s41422-020-00432-2 (2020).

33 Liebschner, D. et al. Macromolecular structure determination using X-rays, neutrons and electrons: recent developments in Phenix. *Acta crystallographica. Section D*, Structural biology 75, 861–877, doi:10.1107/S2059798319011471 (2019).

34 Roh, S.-H. et al. Cryo-EM and MD infer water-mediated proton transport and autoinhibition mechanisms of V(o) complex. Science advances 6, eabb9605, doi:10.1126/sciadv.abb9605 (2020).

35 Chandler, D. Interfaces and the driving force of hydrophobic assembly. Nature 437, 640–647, doi:10.1038/nature04162 (2005).

36 Tanford, C. How protein chemists learned about the hydrophobic factor. Protein science : a publication of the Protein Society 6, 1358–1366, doi:10.1002/pro.5560060627 (1997).

37 Rasaiah, J. C., Garde, S. & Hummer, G. Water in Nonpolar Confinement: From Nanotubes to Proteins and Beyond. Annual Review of Physical Chemistry 59, 713–740, doi:10.1146/annurev.physchem.59.032607.093815 (2008).

38 Striolo, A. Interfacial water studies and their relevance for the energy sector. Molecular Physics 114, 2615–2626, doi:10.1080/00268976.2016.1237685 (2016).

39 Heidari, M., Kremer, K., Potestio, R. & Cortes-Huerto, R. Finite-size integral equations in the theory of liquids and the thermodynamic limit in computer simulations. Molecular Physics 116, 3301–3310, doi:10.1080/00268976.2018.1482429 (2018).

40 Zhang, B. W., Cui, D., Matubayasi, N. & Levy, R. M. The Excess Chemical Potential of Water at the Interface with a Protein from End Point Simulations. The journal of physical chemistry. B 122, 4700–4707, doi:10.1021/acs.jpcb.8b02666 (2018).

41 Weber, V. & Asthagiri, D. Communication: Thermodynamics of water modeled using *ab initio* simulations. The Journal of Chemical Physics 133, doi:10.1063/1.3499315 (2010).

42 Rahbari, A., Hens, R., Dubbeldam, D. & Vlugt, T. J. H. Improving the accuracy of computing chemical potentials in CFCMC simulations. Molecular Physics 117, 3493–3508, doi:10.1080/00268976.2019.1631497 (2019).

43 Hamdi, F. et al. 2.7 Å cryo-EM structure of vitrified M. musculus H-chain apoferritin from a compact 200 keV cryo-microscope. PLoS One 15, e0232540–e0232540, doi:10.1371/journal.pone.0232540 (2020).

44 Yip, K. M., Fischer, N., Paknia, E., Chari, A. & Stark, H. Atomic-resolution protein structure determination by cryo-EM. Nature 587, 157–161, doi:10.1038/s41586-020-2833-4 (2020).

45 Nakane, T. et al. Single-particle cryo-EM at atomic resolution. Nature 587, 152–156, doi:10.1038/s41586-020-2829-0 (2020).

46 Nazarov, S., Myasnikov, A., Stahlberg, H. Cryo-EM structure of mouse heavy-chain apoferritin. To be published, 10.2210/pdb8RQB/pdb (2024).

47 Maki-Yonekura, S., Kawakami, K., Takaba, K., Hamaguchi, T. & Yonekura, K. Measurement of charges and chemical bonding in a cryo-EM structure. Commun Chem 6, 98, doi:10.1038/s42004-023-00900-x (2023).

48 Nakane, T. et al. Single-particle cryo-EM at atomic resolution. Nature 587, 152–156, doi:10.1038/s41586-020-2829-0 (2020).

49 Berman, H. M. et al. The Protein Data Bank. Nucleic Acids Res 28, 235–242, doi:10.1093/nar/28.1.235 (2000).

50 Merzel, F. & Smith, J. C. Is the first hydration shell of lysozyme of higher density than bulk water? Proceedings of the National Academy of Sciences 99, 5378–5383, doi:doi:10.1073/pnas.082335099 (2002).

51 Iudin, A. et al. EMPIAR: the Electron Microscopy Public Image Archive. Nucleic Acids Res 51, D1503–D1511, doi:10.1093/nar/gkac1062 (2023).

52 Carugo, O. & Bordo, D. How many water molecules can be detected by protein crystallography? Acta Crystallogr D Biol Crystallogr 55, 479–483, doi:10.1107/s0907444998012086 (1999).

53 Levy, Y. & Onuchic, J. N. Water and proteins: A love–hate relationship. Proceedings of the National Academy of Sciences 101, 3325–3326, doi:doi:10.1073/pnas.0400157101 (2004).

54 Kriegel, M. & Muller, Y. A. De novo prediction of explicit water molecule positions by a novel algorithm within the protein design software MUMBO. Scientific Reports 13, 16680, doi:10.1038/s41598-023-43659-w (2023).

55 Rudling, A., Orro, A. & Carlsson, J. Prediction of Ordered Water Molecules in Protein Binding Sites from Molecular Dynamics Simulations: The Impact of Ligand Binding on Hydration Networks. Journal of Chemical Information and Modeling 58, 350–361, doi:10.1021/acs.jcim.7b00520 (2018).

56 Berkeley, R. F., Cook, B. D. & Herzik, M. A., Jr. Machine learning approaches to cryoEM density modification differentially affect biomacromolecule and ligand density quality. Front Mol Biosci 11, 1404885, doi:10.3389/fmolb.2024.1404885 (2024).

57 Pronk, S. et al. GROMACS 4.5: a high-throughput and highly parallel open source molecular simulation toolkit. Bioinformatics 29, 845–854, doi:10.1093/bioinformatics/btt055 (2013).

58 Maier, J. A. et al. ff14SB: Improving the Accuracy of Protein Side Chain and Backbone Parameters from ff99SB. Journal of Chemical Theory and Computation 11, 3696–3713, doi:10.1021/acs.jctc.5b00255 (2015).

59 Jorgensen, W. L., Chandrasekhar, J., Madura, J. D., Impey, R. W. & Klein, M. L. Comparison of simple potential functions for simulating liquid water. The Journal of Chemical Physics 79, 926–935, doi:10.1063/1.445869 (1983).

60 Berendsen, H. J. C., Postma, J. P. M., van Gunsteren, W. F., DiNola, A. & Haak, J. R. Molecular dynamics with coupling to an external bath. The Journal of Chemical Physics 81, 3684–3690, doi:10.1063/1.448118 (1984).

61 Parrinello, M. a. R., A. Polymorphic transitions in single crystals: A new molecular dynamics method. Journal of Applied Physics 52, 7182–7190 (1981).

62 Essmann, U. et al. A Smooth Particle Mesh Ewald Method. Journal of Chemical Physics 103, 8577–8593, doi:Doi 10.1063/1.470117 (1995).

63 Zivanov, J., Nakane, T. & Scheres, S. H. W. Estimation of high-order aberrations and anisotropic magnification from cryo-EM data sets in RELION-3.1. IUCrJ 7, 253–267, doi:10.1107/S2052252520000081 (2020).

64 Punjani, A., Rubinstein, J. L., Fleet, D. J. & Brubaker, M. A. cryoSPARC: algorithms for rapid unsupervised cryo-EM structure determination. Nat Methods 14, 290–296, doi:10.1038/nmeth.4169 (2017).

65 Grant, T. & Grigorieff, N. Measuring the optimal exposure for single particle cryo-EM using a 2.6 A reconstruction of rotavirus VP6. eLife 4, e06980, doi:10.7554/eLife.06980 (2015).

66 Zivanov, J., Nakane, T. & Scheres, S. H. W. A Bayesian approach to beam-induced motion correction in cryo-EM single-particle analysis. IUCrJ 6, 5–17, doi:10.1107/S205225251801463X (2019).

67 Beckers, M., Jakobi, A. J. & Sachse, C. Thresholding of cryo-EM density maps by false discovery rate control. IUCrJ 6, 18–33, doi:10.1107/S2052252518014434 (2019).

68 Beckers, M., Palmer, C. M. & Sachse, C. Confidence maps: statistical inference of cryo-EM maps. Acta Crystallogr D Struct Biol 76, 332–339, doi:10.1107/S2059798320002995 (2020).

69 Goddard, T. D. et al. UCSF ChimeraX: Meeting modern challenges in visualization and analysis. Protein Sci 27, 14–25, doi:10.1002/pro.3235 (2018).

70 Pettersen, E. F. et al. UCSF Chimera--a visualization system for exploratory research and analysis. J Comput Chem 25, 1605–1612, doi:10.1002/jcc.20084 (2004).

