## supplemental tables and figures for "De Novo Hydration of Cryo-EM Reconstructions through Molecular Dynamics Simulations of the Excess Chemical Potential"

**Table S1** Summary information of 9 cryo-EM structures of apoferritin (Group A) with resolution higher than 1.5 Å and 17 cryo-EM structures of apoferritin (Group B) with resolution between 1.5 and 2.0 Å.

| PDB ID | Year of Deposition | Resolution (Å) | Refinement Software | Number of Cryo-EM Water per Chain |
|---|---|---|---|---|
| **Group A (9)** | | | | |
| 8RQB | 2024 | 1.09 | Unknown | 110 |
| 7A6A | 2020 | 1.15 | Refmac | 194 |
| 8J5A | 2023 | 1.19 | Refmac | 145 |
| 7A4M | 2020 | 1.22 | Refmac | 110 |
| 6Z6U | 2020 | 1.25 | Refmac | 139 |
| 7RRP | 2021 | 1.27 | Phenix | 102 |
| 7A6B | 2020 | 1.33 | Refmac | 162 |
| 7K3V | 2020 | 1.34 | Phenix | 111 |
| 7K3W | 2020 | 1.36 | Phenix | 111 |
| **Group B (17)** | | | | |
| 6Z9E | 2020 | 1.55 | Refmac | 146 |
| 6Z9F | 2020 | 1.56 | Refmac | 172 |
| 7R5O | 2022 | 1.58 | Refmac | 139 |
| 7KOD | 2020 | 1.65 | Phenix | 73 |
| 8EMQ | 2022 | 1.66 | Phenix | 124 |
| 8EN7 | 2022 | 1.68 | Phenix | 106 |
| 6V21 | 2019 | 1.75 | Phenix | 75 |
| 8CPU | 2023 | 1.76 | Phenix | 73 |
| 8CPV | 2023 | 1.76 | Phenix | 71 |
| 8CPX | 2023 | 1.76 | Phenix | 70 |
| 7ZG7 | 2022 | 1.77 | Phenix | 111 |
| 8CPT | 2023 | 1.79 | Phenix | 79 |
| 8CPW | 2023 | 1.79 | Phenix | 78 |
| 8CPM | 2023 | 1.81 | Phenix | 81 |
| 8CPS | 2023 | 1.82 | Phenix | 84 |
| 6S61 | 2019 | 1.84 | Phenix | 160 |
| 7VD8 | 2021 | 1.96 | Phenix | 103 |

**Table S2** The observation of high- and low-consensus waters in cryo-EM maps across various resolutions (1.34 Å, 1.49 Å, ... 3.84 Å) and at three different false discovery rate (FDR) thresholds: 0.1%, 1%, and 5%.

| Water Confidence | Resolution (Å) | | | | | | | | | |
|---|---|---|---|---|---|---|---|---|---|---|
| | 1.34 | 1.49 | 1.53 | 1.6 | 1.67 | 1.82 | 2.21 | 2.68 | 2.92 | 3.84 |
| **lower** | No | No | No at 0.1%, partial overlap at 1 and 5% | No at 0.1 and 1%, partial at 5% | No at 0.1%, partial overlap at 1 and 5% | No | No | No | No | No |
| **lower** | No at 0.1 and 1%, partial overlap at 5% | No at 0.1%, partial overlap at 1 and 5% | No | No | No | No | Noisy and partial at 0.1%, noisy at 1 and 5% | Noisy and partial | No | No |
| **lower** | Yes | Partial | Partial at 0.1%, complet e at 1 and 5% | No at 0.1%, partial at 1%, complet e overlap at 5% | Partial | No at 0.1%, partial overlap at 1 and 5% | No | Partial at 0.1%, noisy at 1 and 5% | Noisy and partial at 0.1%, noisy at 1 and 5% | No |
| **lower** | No at 0.1%, partial at 1 and 5% | No at 0.1 and 1%, partial overlap at 5% | No | No at 0.1%, partial at 1 and 5% | No | No at 0.1%, partial overlap at 1 and 5% | No at 0.1 and 1%, partial overlap at 5% | No | No | No |
| **lower** | No | No | No | No | No | No | No | No | No | No |
| **lower** | Partial | Partial | No at 0.1%, partial overlap at 1 and 5% | No at 0.1%, partial at 1 and 5% | No at 0.1%, partial overlap at 1 and 5% | No at 0.1 and 1%, partial overlap at 5% | No at 0.1%, partial and noisy at 1 and 5% | No | No | No |
| **lower** | No at 0.1 and 1%, partial overlap at 5% | No at 0.1%, partial overlap at 1 and 5% | No | No at 0.1%, partial at 1 and 5% | No at 0.1 and 1%, partial overlap at 5% | Partial | No at 0.1%, partial overlap at 1 and 5% | No | No | No |
| **lower** | No at 0.1%, partial at 1 and 5% | No | No | No | No | No at 0.1%, partial overlap at 1 and 5% | Noisy and partial at 0.1%, noisy at 1 and 5% | Noisy and partial at 0.1 and 1%, noisy at 5% | No | No |
| **lower** | Partial | Partial | No at 0.1%, partial overlap at 1 and 5% | No at 0.1 and 1%, noisy at 5% | No | No at 0.1%, partial overlap at 1 and 5% | No | No | No | No |
| **lower** | Partial | Partial | No at 0.1 and 1%, partial at 5% | No at 0.1 and 1, noisy at 5% | No | No at 0.1%, partial overlap at 1 and 5% | No | No | No | No |
| **lower** | Yes | Partial | Yes | Yes | Yes | Yes | Yes | Noisy | No at 0.1% and 1% and partial | No |

| | | | | | | | | | | |
|---|---|---|---|---|---|---|---|---|---|---|
| | | | | | | | | | overlap at 5% | |
| **lower** | Yes | Yes | Yes | Yes | Yes | Yes | Noisy | Noisy | No | No |
| **lower** | Yes | Yes | Yes | No at 0.1, noisy at 1 and 5% | Yes | Yes | Yes | No at 0.1, noisy and partial overlap at 1% and 5% | No | No |
| **lower** | Noisy | Noisy | Yes | Yes | Yes | Yes | Noisy | Noisy and partial | No | No |
| **lower** | Yes | Yes | Noisy | Yes | Yes | Partial at 0.1%, complete at 1 and 5% | Yes | Yes | Yes | No |
| **lower** | Yes | Partial at 0.1 and 1%, complete overlap at 5% | No at 0.1%, partial overlap at 1 and 5% | Yes | No at 0.1%, partial overlap at 1 and 5% | Partial | Yes | Partial at 0.1 and 1%, noisy at 5% | No | No |
| **lower** | Partial at 0.1%, complete overlap at 1 and 5% | Yes | Partial at 0.1 and 1, complete overlap at 5% | Partial at 0.1 and 1%, noisy at 5% | Partial at 0.1, complete overlap at 1 and 5% | Yes | Yes | No at 0.1%, partial at 1 and 5% | No | No |
| **lower** | Partial at 0.1%, complete overlap at 1 and 5% | Partial | Partial | No | No | No | No at 0.1%, partial and noisy at 1 and 5% | Noisy | No | No |
| **lower** | Yes | Yes and elongated density | No at 0.1%, overlap at 1 and 5% with elongated density | Yes | No at 0.1%, partial overlap at 1 and 5% | Partial | Noisy | Partial at 0.1 and 1%, partial and noisy at 5% | No | No |
| **lower** | Partial at 0.1 and 1%, complete overlap at 5% | Yes | Partial at 0.1 and 1, complete overlap at 5% | No | No | No at 0.1%, partial overlap at 1 and 5% | Noisy | Noisy | No | No |
| **lower** | Yes | Yes | Partial at 0.1, complete overlap at 1 and 5% | Yes | Partial at 0.1, complete overlap at 1 and 5% | Yes | Noisy | No | No | No |
| **lower** | Yes | Yes | Yes | Partial | Partial overlap at 0.1%, complete overlap 1 and 5% | Yes | Yes | No | No | No |

| | | | | | | | | | |
|---|---|---|---|---|---|---|---|---|---|
| **lower** | Yes | Partial at 0.1%, complete overlap at 1 and 5% | Yes | Partial at 0.1%, complete overlap at 1 and 5% | Partial | No at 0.1 and 1%, partial overlap at 5% | Noisy | Noisy | No | No |
| **lower** | Partial at 0.1%, noisy and partial at 1%, noisy at 5% | Yes | Yes | No | No | No at 0.1%, partial overlap at 1 and 5% | Noisy | No at 0.1%, noisy and partial at 1 and 5% | No | No |
| **high** | Yes | Yes | Yes | Yes | Yes | Yes | Yes | Noisy | No | No |
| **high** | Yes | Yes | Yes | Yes | Yes | Partial overlap at 0.1 and 1%, complete overlap at 5% | Noisy | Yes | No | No |
| **high** | Yes | Yes | Yes | Yes | Yes | Yes | Noisy | Yes | No | No |
| **high** | Yes | Yes | Yes | Yes | Yes | Yes | Yes | Noisy | Partial at 0.1%, noisy at 1 and 5% | No |
| **high** | Yes | Yes | Yes | Yes | Yes | Yes | Noisy | Partial at 0.1, noisy at 1 and 5% | No at 0.1%, noisy at 1 and 5% | No |
| **high** | Yes | Yes | Yes | Yes | Partial overlap at 0.1 and 1%, complete overlap at 5% | Partial overlap at 0.1 and 1%, complete overlap at 5% | Noisy | Noisy | No | No |
| **high** | Yes | Yes | Yes | Yes | Yes | Yes | Yes | Noisy | No at 0.1%, partial overlap at 1% and 5% | No |
| **high** | Yes | Yes | Yes | Yes | Partial at 0.1, complete overlap at 1 and 5% | Yes | Noisy | Partial at 0.1, complete overlap at 1, noisy at 5% | No | No |
| **high** | Yes | Yes | Yes | Yes | Yes | Yes | Yes | Partial at 0.1, complete overlap at 1 and 5% | No | No |
| **high** | Yes | Yes | Yes | Yes | Yes | Yes | Yes | Noisy | No at 0.1%, partial overlap | No |

| | | | | | | | | | | |
|---|---|---|---|---|---|---|---|---|---|---|
| | | | | | | | | | at 1% and 5% | |
| **high** | Yes | Yes | Yes | Yes | Yes | Yes | Yes | Noisy | No | No |
| **high** | Yes | Yes | Yes | Yes | Noisy | Noisy | No | No at 0.1, noisy and partial overlap at 1% and 5% | No at 0.1%, noisy at 1 and 5% | No |
| **high** | Yes | Yes | Yes | Yes | Yes | Yes | Yes | Noisy | No | No |
| **high** | Yes | Yes | Yes | Yes | Yes | Yes | Noisy | Noisy | Noisy | No |
| **high** | Noisy | Yes | Yes | Yes | Yes | Yes | Noisy | Noisy and partial at 0.1 and 1%, noisy at 5% | No | No |
| **high** | Yes | Yes | Yes | Yes | Yes | Yes | Yes | Partial at 0.1 and 1%, noisy at 5% | No | No |
| **high** | Yes | Yes | Yes | Yes | Yes | Yes | Noisy | Noisy | No | No |
| **high** | Yes | Yes | Yes | Yes | Yes | Yes | Noisy | Noisy and partial at 0.1 and 1%, noisy at 5% | No | No |
| **high** | Yes | Yes | Yes | Yes | Yes | Yes | Noisy | Partial at 0.1%, complet e overlap at 1 and 5% | No at 0.1%, partial overlap at 1% and 5% | No |
| **high** | Yes | Yes | Yes | Yes | Yes | Yes | Noisy | Noisy | No | No |
| **high** | Yes | Yes | Yes | Yes | Yes | Yes | Noisy | Noisy | No at 0.1 and 1%, noisy at 5% | No |
| **high** | Yes | Yes | Yes | Yes | Yes | Yes | Yes | Yes | No | No |
| **high** | Yes | Yes | Yes | Yes | Yes | Yes | Yes | No at 0.1, noisy and partial overlap at 1% and 5% | No at 0.1, noisy and partial overlap at 1% and 5% | No |
| **high** | Yes | Yes | Yes | Yes | Yes | Yes | Noisy | Yes | No at 0.1%, partial at 1 and 5% | No |
| **high** | Yes | Yes | Yes | Yes | Yes | Yes | Noisy | Noisy | Noisy | No |
| **high** | Yes | Yes | Yes | Yes | Yes | Yes | Noisy | Yes | Partial at 0.1, complet e | No |

| | | | | | | | | | | |
|---|---|---|---|---|---|---|---|---|---|---|
| | | | | | | | | | overlap at 1 and 5% | |
| **high** | Yes | Yes | Yes | Yes | Yes | Yes | Yes | Yes | No at 0.1%, partial overlap at 1% and noisy at 5% | No |
| **high** | Yes | Yes | Yes | Yes | Yes | Yes | Noisy | Noisy | No at 0.1 and 1%, noisy at 5% | No |
| **high** | Yes | Yes | Yes | Yes | Yes | Yes | Noisy | Yes | Noisy | No |
| **high** | Yes | Yes | Yes | Yes | Yes | Yes | Noisy | Noisy | No | No |
| **high** | Yes | Yes | Yes | Yes | Yes | Yes | Noisy | Noisy | No at 0.1%, partial at 1 and 5% | No |
| **high** | Yes | Yes | Yes | Yes | Yes | Yes | Noisy | Noisy | No | No |
| **high** | Yes | Yes | Yes | Yes | Yes | Yes | Noisy | Noisy | No | No |
| **high** | Yes | Yes | Yes | Yes | Yes | Yes | Yes | Yes | No | No |
| **high** | Yes | Yes | Yes | Yes | Yes | Yes | Noisy | Noisy | No at 0.1% and 1%, noisy and partial overlap at 5% | No |
| **high** | Yes | Yes | Yes | Yes | Yes | Yes | Noisy | Yes | No | No |
| **high** | Yes | Yes | Yes | Yes | Yes | Yes | Yes | Partial | No | No |
| **high** | Yes | Yes | Yes | Yes | Yes | Yes | Yes | Noisy | No | No |
| **high** | Yes | Yes | Yes | Yes | Yes | Yes | Yes | No at 0.1, noisy and partial overlap at 1% and 5% | No | No |
| **high** | Yes | Yes | Yes | Yes | Yes | Yes | Noisy | Partial at 0.1%, noisy at 1 and 5% | No | No |
| **high** | Yes | Yes | Yes | Yes | Yes | Yes | Noisy | Noisy | No | No |
| **high** | Yes | Yes | Yes | Yes | Yes | Partial at 0.1 and 1%, complet e at 5% | Yes | Noisy and partial at 0.1%, noisy at 1 and 5% | No | No |
| **high** | Yes | Yes | Yes | Yes | Yes | Yes | Yes | Noisy | Yes | No |
| **high** | Yes | Yes | Yes | Yes | Yes | Noisy | Noisy | Noisy | No | No |
| **high** | Yes | Yes | Yes | Yes | Yes | Yes | Noisy | Noisy | No | No |

| | | | | | | | | | | |
|---|---|---|---|---|---|---|---|---|---|---|
| **high** | Yes | Yes | Yes | Yes | Yes | Yes | Noisy | Yes | No | No |
| **high** | Yes | Yes | Yes | Yes | Yes | Partial at 0.1 and 1%, complet e at 5% | Noisy | Yes | No | No |
| **high** | Yes | Yes | Yes | Yes | Yes | Yes | Noisy | Partial at 0.1%, complet e overlap at 1 and 5% | No | No |
| **high** | Yes | Yes | Yes | Yes | Yes | Yes | Noisy | Noisy | Partial and noisy | No |
| **high** | Yes | Yes | Yes | Yes | Yes | Yes | Noisy | Partial at 0.1 and 1%, complet e overlap at 5% | No | No |
| **high** | Yes | Yes | Yes | Yes | Yes | Yes | Yes | Noisy | No at 0.1%, partial at 1% and complet e at 5% | No |
| **high** | Yes | Yes | Yes | Yes | Yes | Yes | Yes | Yes | No | No |
| **high** | Yes | Yes | Yes | Yes | Yes | Yes | Noisy | Yes | Noisy | No |

**Figure S1:** Final Apoferritin maps are shown at different resolutions (1.34 Å, 1.49 Å, ... 3.84 Å), with Y168 displayed at the bottom of each respective map at a 1% FDR threshold.

**Figure S2:** The plot showing the number of water molecules per Apoferritin residue as a function of map resolution is based on a 0.1% (**A**) and 5% (**B**) false discovery rate (FDR) threshold.

**Figure S3:** The interaction energy is plotted against the distance for two hydrogen-bonded water molecules(shown in inset) in gas phase using TIP3P model. The curves for Coulombic energy, Lennard-Jones energy and the total interaction energy are shown in grey, blue and red respectively.

**Figure S4:** The histogram illustrates the distribution of distances between matched predicted waters and experimental waters with a 2.2 Å distance cutoff.

#### SI Figure S1

Final apoferritin maps at different resolutions along with the Y168 displayed at the bottom of the respective map

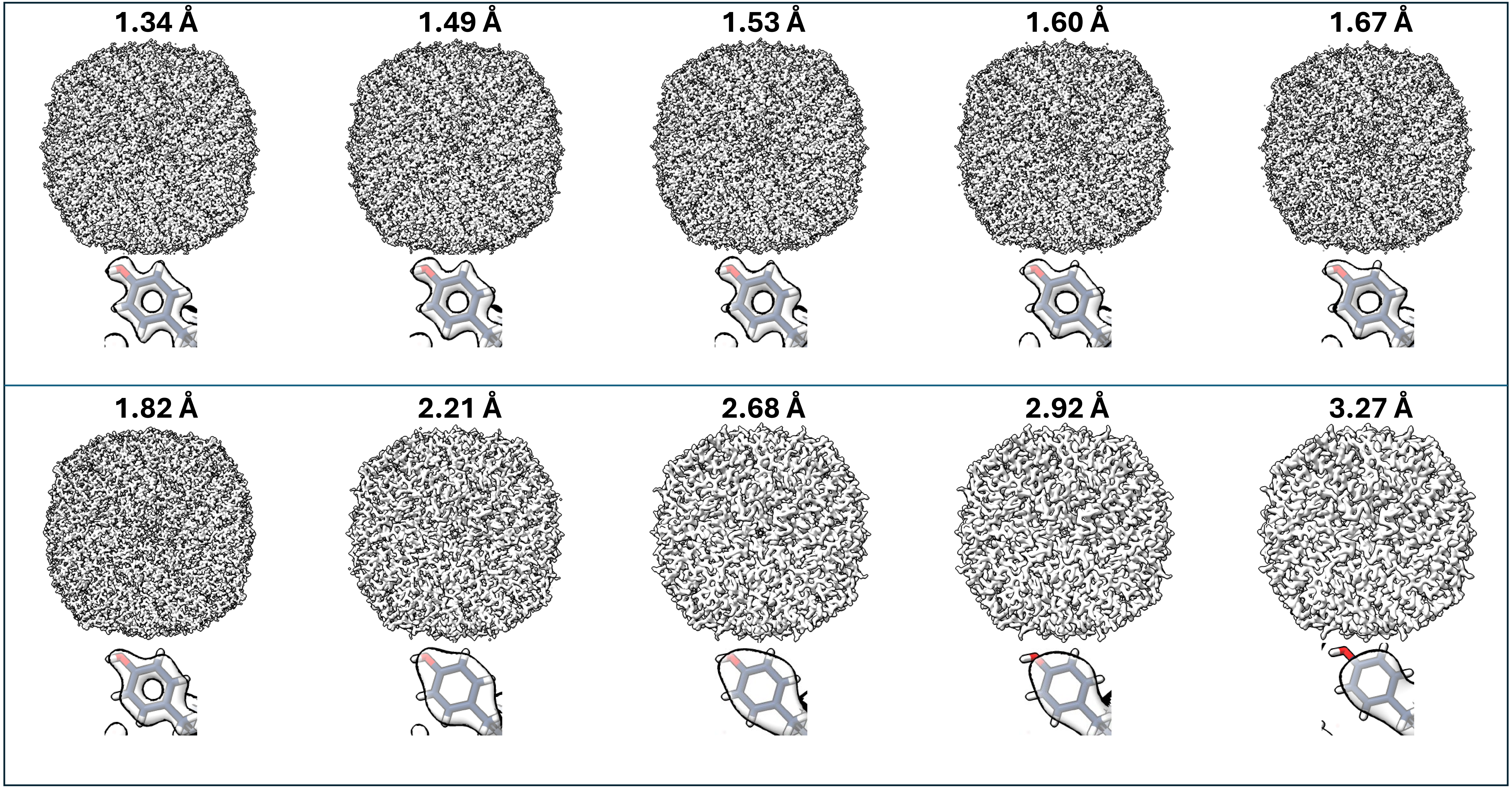

SI Figure S2

The number of water molecules per apoferritin residue as a function of map resolution at 0.1% and 5% FDR threshold

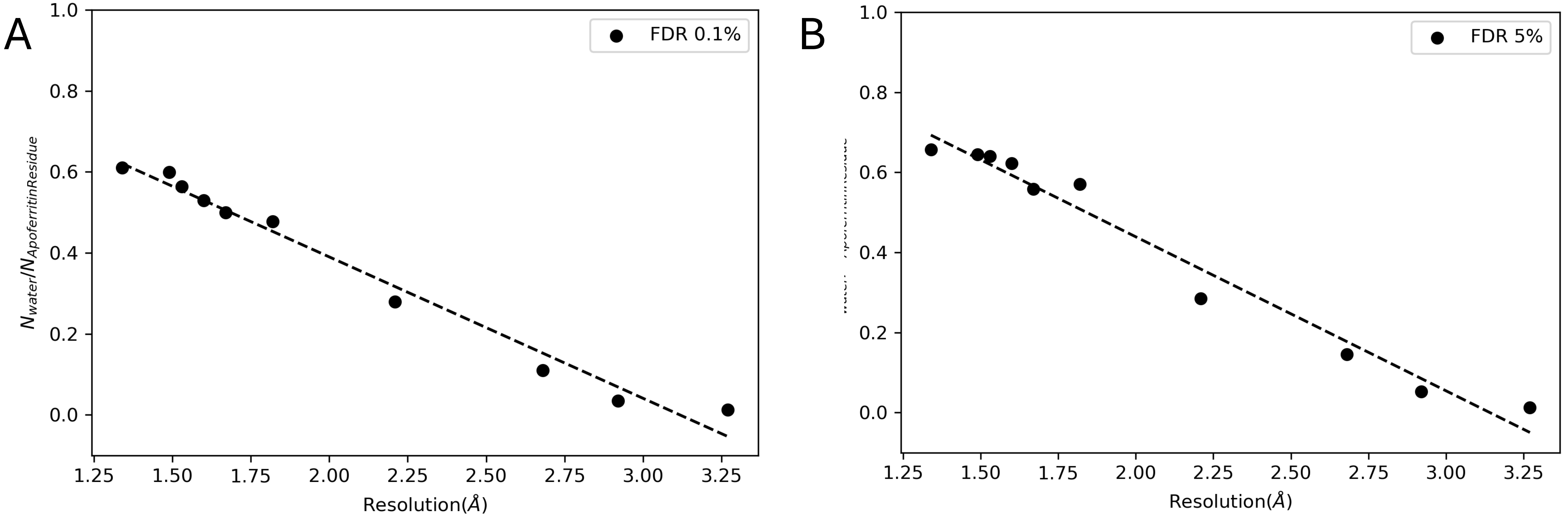

SI Figure S3

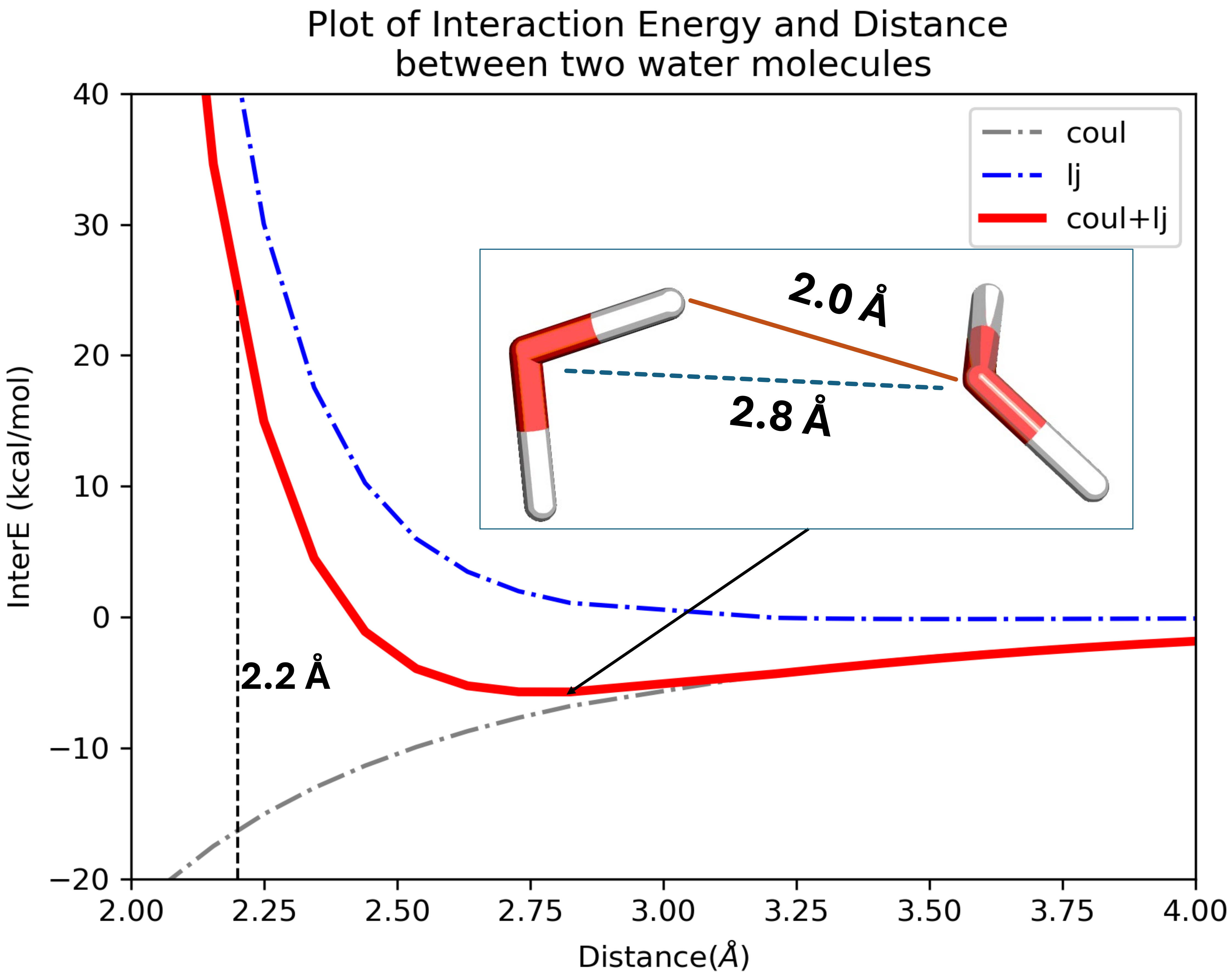

Plot of Interaction Energy and Distance
between two water molecules
coul
lj
coul+lj
2.0 Å
2.8 Å
2.2 Å
InterE (kcal/mol)
Distance(Å)
40
30
20
10
0
−10
−20
2.00
2.25
2.50
2.75
3.00
3.25
3.50
3.75
4.00

### SI Figure S4

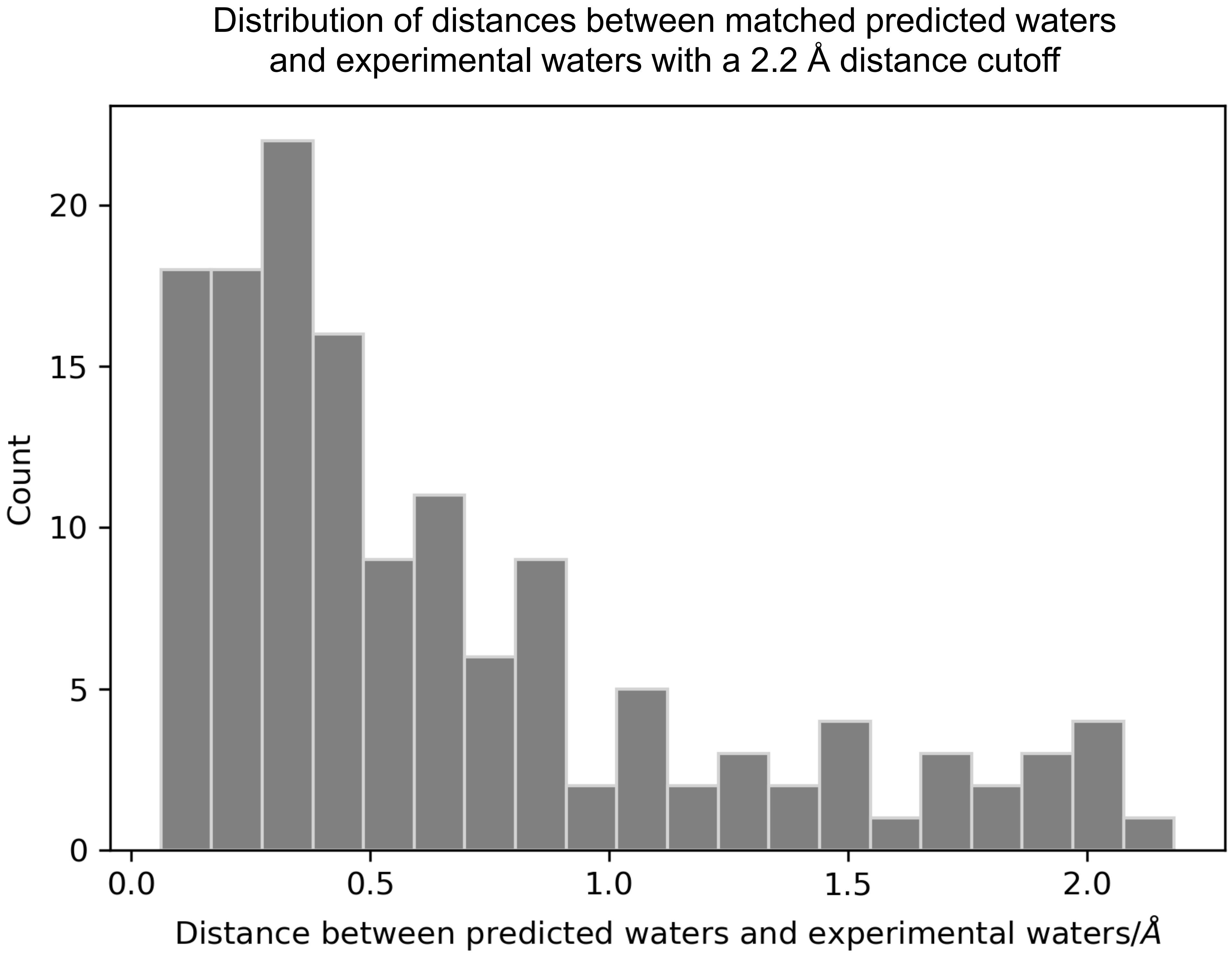
Distribution of distances between matched predicted waters
and experimental waters with a 2.2 Å distance cutoff
Count
20
15
10
5
0
0.0
0.5
1.0
1.5
2.0
Distance between predicted waters and experimental waters/Å
